# Shedding Light on Microbial Dark Matter with A Universal Language of Life

**DOI:** 10.1101/2020.12.23.424215

**Authors:** A Hoarfrost, A Aptekmann, G Farfañuk, Y Bromberg

## Abstract

The majority of microbial genomes have yet to be cultured, and most proteins predicted from microbial genomes or sequenced from the environment cannot be functionally annotated. As a result, current computational approaches to describe microbial systems rely on incomplete reference databases that cannot adequately capture the full functional diversity of the microbial tree of life, limiting our ability to model high-level features of biological sequences. The scientific community needs a means to capture the functionally and evolutionarily relevant features underlying biology, independent of our incomplete reference databases. Such a model can form the basis for transfer learning tasks, enabling downstream applications in environmental microbiology, medicine, and bioengineering. Here we present LookingGlass, a deep learning model capturing a “universal language of life”. LookingGlass encodes contextually-aware, functionally and evolutionarily relevant representations of short DNA reads, distinguishing reads of disparate function, homology, and environmental origin. We demonstrate the ability of LookingGlass to be fine-tuned to perform a range of diverse tasks: to identify novel oxidoreductases, to predict enzyme optimal temperature, and to recognize the reading frames of DNA sequence fragments. LookingGlass is the first contextually-aware, general purpose pre-trained “biological language” representation model for short-read DNA sequences. LookingGlass enables functionally relevant representations of otherwise unknown and unannotated sequences, shedding light on the microbial dark matter that dominates life on Earth.

**Availability:** The pretrained LookingGlass model and the transfer learning-derived models demonstrated in this paper are available in the LookingGlass release v1.0^1^. The open source *fastBio* Github repository and python package provides classes and functions for training and fine tuning deep learning models with biological data^2^. Code for reproducing analyses presented in this paper are available as an open source Github repository^3^.

## Introduction

The microbial world is dominated by “microbial dark matter” – the majority of microbial genomes remain to be sequenced^4,5^, while the molecular functions of many genes in microbial genomes are unknown^6^. In microbial communities (microbiomes), the combination of these factors compounds this limitation. While the rate of biological sequencing outpaces Moore’s law^7^, our traditional experimental means of annotating these sequences cannot keep pace. Scientists thus typically rely on reference databases which reflect only a tiny fraction of the biological diversity on Earth.

Our reliance on this incomplete annotation of biological sequences propagates significant observational bias toward annotated genes and cultured genomes in describing microbial systems. To break out of this cycle, the scientific community needs a means of representing biological sequences that captures their functional and evolutionary relevance and that is independent of our limited references.

Deep learning is particularly good at capturing complex, high dimensional systems, and is a promising tool for biology^8^. However, deep learning generally requires massive amounts of data to perform well. Meanwhile, collection and experimental annotation of samples is typically time consuming and expensive, and the creation of massive datasets for one study is rarely feasible. The scientific community needs a means of building computational models which can capture biological complexity while compensating for the low sample size and high dimensionality that characterize biology.

Transfer learning provides a solution to the high-dimensionality, low-sample-size conundrum. Transfer learning^9,10^ leverages domain knowledge learned by a model in one training setting and applies it to a different but related problem. This approach is effective because a model trained on a massive amount of data from a particular data modality of interest (e.g. biological sequences) will learn features *general* to that modality in addition to the *specific* features of its learning task. This general pretrained model can then be further trained, or “fine-tuned”, to predict a downstream task of interest more accurately, using less task-specific data, and in shorter training time than would otherwise be possible. In computer vision, for example, by starting from a pretrained model trained on many images, a model of interest doesn’t relearn general image features such as a curve or a corner^11^, but instead can devote its limited dataset to refining the specific parameters of the target task. In natural language processing, a generic language representation model^12^ has been widely applied to diverse text classification tasks, including biomedical text classification^13,14^.

Pretrained models lower the barrier for widespread academic and private sector applications, which typically have small amounts of data and limited computational resources to model relatively complex data. Natural language processing for text, and language modelling in particular, is analogous to biological sequences, in that nucleotides are not independent or identically distributed^15^ and the nucleotide *context* is important for defining the functional role and evolutionary history of the whole sequence.

In genomics and metagenomics, there is no analogous contextually-aware pretrained model that can be generally applied for transfer learning on read-length biological sequences. Some previous studies have obtained important results using transfer learning^16,17^, but were either limited to relatively small training sets for pretraining a model on a closely related prediction task^16^, or relied on gene counts from the relatively well-annotated human genome to compile their training data^17^. Previous works in learning continuous representations of biological sequences^18,19^ and genomes^20^ do not account for the order in which sequences or proteins appear and are thus not contextually-aware. Recent advances in full-length protein sequence representation learning^21–24^ show the potential of a self-supervised learning approach that accounts for sequence context, but these rely on full length protein sequences (ca. 1,000 amino acids or 3,000 nucleotides). Full-length protein sequences are computationally difficult (and sometimes impossible) to assemble from metagenomes, which can produce hundreds of millions of short-read DNA sequences (ca. 60-300 nucleotides) per sample. To capture the full functional diversity of the microbial world, we need a contextually-relevant means to represent the functional and evolutionary features of biological sequences from microbial communities, in the short, fragmented form in which they are sampled from their environment.

A biological ‘universal language of life’ should reflect functionally and evolutionarily relevant features that underly biology as a whole and facilitate diverse downstream transfer learning tasks. Here, we present LookingGlass, a biological language model and sequence encoder, which produces contextually relevant embeddings for any biological sequence across the microbial tree of life. LookingGlass is trained and optimized for read-length sequences, such as those produced by the most widely used sequencing technologies^25^. For metagenomes in particular, a read-level model avoids the need for assembly, which has a high computational burden and potential for error. We also focus on Bacterial and Archaeal sequences, although we include a discussion of the possibility for Eukaryotic and human-specific models below.

We demonstrate the functional and evolutionary relevance of the embeddings produced by LookingGlass, and its broad utility across multiple transfer learning tasks relevant to functional metagenomics. LookingGlass produces embeddings that differentiate sequences with different molecular functions; identifies homologous sequences, even at low sequence similarities where traditional bioinformatics approaches fail; and differentiates sequences from disparate environmental contexts. Using transfer learning, we demonstrate how LookingGlass can be used to illuminate the “microbial dark matter” that dominates environmental settings by developing an ‘oxidoreductase classifier’ that can identify novel oxidoreductases (enzymes responsible for electron transfer, and the basis of all metabolism) with very low sequence similarity to those seen during training. We also demonstrate LookingGlass’ ability to predict enzyme optimal temperatures from short-read DNA fragments; and to recognize the reading frame (and thus “true” amino acid sequence) encoded in short-read DNA sequences with high accuracy.

The transfer learning examples shown here, aside from providing useful models in and of themselves, are intended to show the broad types of questions that can be addressed with transfer learning from a single pretrained model. These downstream models can illuminate the functional role of “microbial dark matter” by leveraging domain knowledge of the functional and evolutionary features underlying microbial diversity as a whole. More generally, LookingGlass is intended to serve as the scientific community’s ‘universal language of life’ that can be used as the starting point for transfer learning in biological applications, and metagenomics in particular.

## Methods

### I. LookingGlass design and optimization

#### Dataset Generation

The taxonomic organization of representative Bacterial and Archaeal genomes was determined from the Genome Taxonomy Database, GTDB^26^ (release 89.0). The complete genome sequences were downloaded via the NCBI Genbank ftp^27^. This resulted in 24,706 genomes, comprising 23,458 Bacterial and 1,248 Archaeal genomes.

Each genome was split into read-length chunks. To determine the distribution of realistic read lengths produced by next-generation short read sequencing machines, we obtained the BioSample IDs^27^ for each genome, where they existed, and downloaded their sequencing metadata from the MetaSeek^28^ database using the MetaSeek API. We excluded samples with average read lengths less than 60 or greater than 300 base pairs. This procedure resulted in 7,909 BioSample IDs. The average read lengths for these sequencing samples produced the ‘read-length distribution’ (SI Fig 1) with a mean read length of 136bp. Each genome was split into read-length chunks (with zero overlap in order to maximize information density and reduce data redundancy in the dataset): a sequence length was randomly selected with replacement from the read-length distribution and a sequence fragment of that length was subset from the genome, with a 50% chance that the reverse complement was used. The next sequence fragment was chosen from the genome starting at the end point of the previous read-length chunk, using a new randomly selected read length, and so on. To ensure that genomes in the training, validation, and test sets had low sequence similarity, the sets were split along taxonomic branches such that genomes from the *Actinomycetales*, *Rhodobacterales*, *Thermoplasmata*, and *Bathyarchaeia* were partitioned into the validation set; genomes from the *Bacteroidales*, *Rhizobiales, Methanosarcinales,* and *Nitrososphaerales* were partitioned into the test set; and the remaining genomes remained in the training set. This resulted in 529,578,444 sequences in the training set, 57,977,217 sequences in the validation set, and 66,185,518 sequences in the test set. We term this set of reads the *GTDB representative* set (Table 1).

**Table 1.**
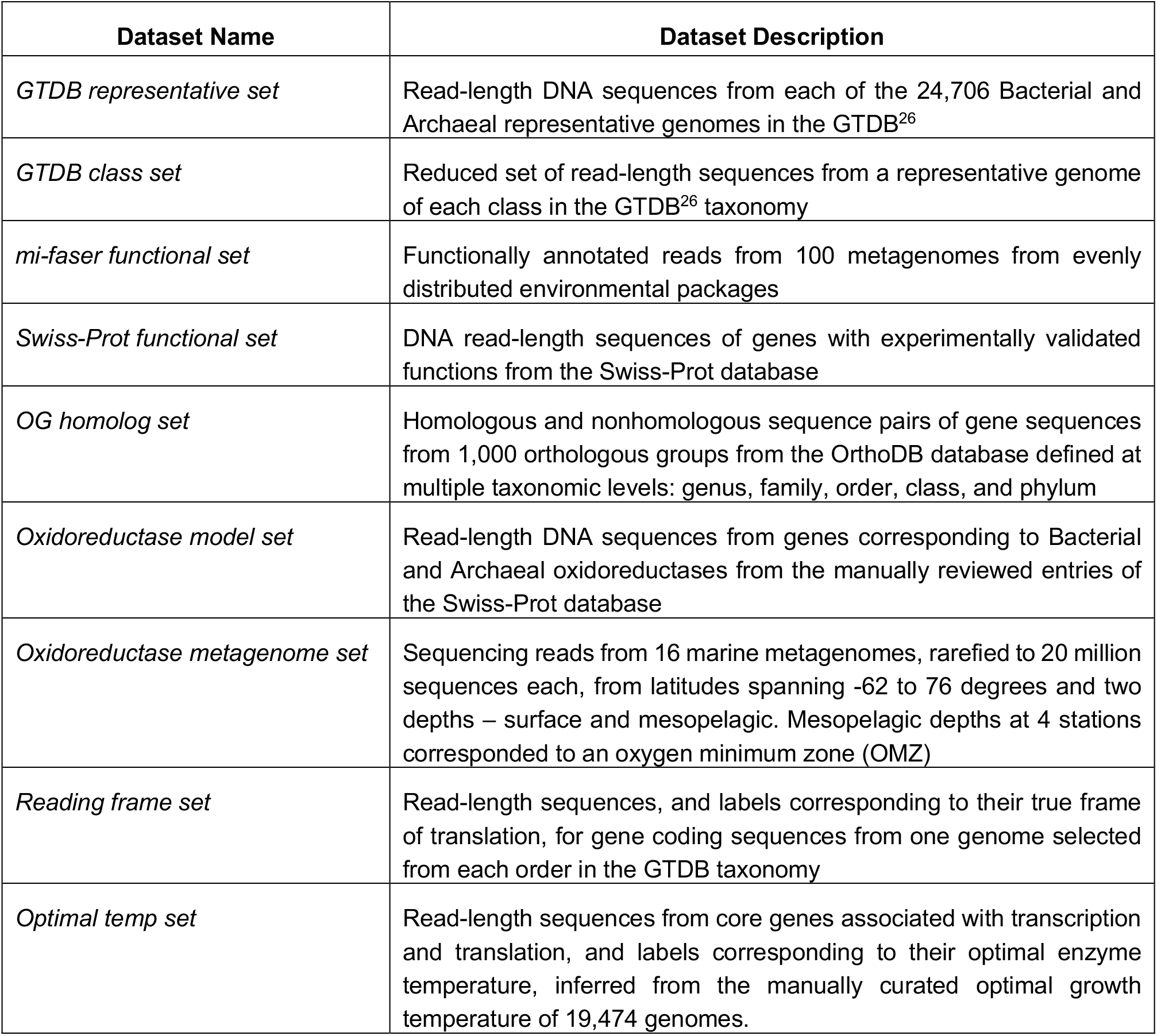
Summary table of datasets used.

The amount of data needed for training was also evaluated (SI Fig 2). Progressively larger amounts of data were tested by selecting at random 1, 10, 100, or 500 read-length chunks from each of the GTDB representative genomes in the *GTDB representative training set*. Additionally, the performance of smaller but more carefully selected datasets, representing the diversity of the microbial tree of life, were tested by selecting for training one genome at random from each taxonomic class or order in the GTDB taxonomy tree. In general, better accuracy was achieved in fewer epochs with a greater amount of sequencing data (SI Fig 2); however, a much smaller amount of data performed better if a representative genome was selected from each GTDB taxonomy class.

The final LookingGlass model was trained on this class-level partition of the microbial tree of life. We term this dataset the *GTDB class set* (Table 1). The training, validation, and test sets were split such that no classes overlapped across sets: the validation set included 8 genomes from each of the classes Actinobacteria, Alphaproteobacteria, Thermoplasmata, and Bathyarchaeia (32 total genomes); the test set included 8 genomes from each of the classes Bacteroidia, Clostridia, Methanosarcinia, and Nitrososphaeria (32 total genomes); and the training set included 1 genome from each of the remaining classes (32 archaeal genomes and 298 bacterial genomes for a total of 330 genomes). This resulted in a total of 6,641,723 read-length sequences in the training set, 949,511 in the validation set, and 632,388 in the test set (SI Table 1).

#### Architecture design and training

Recurrent Neural Networks (RNNs) are a type of neural network designed to take advantage of the context dependence of sequential data (such as text, video, audio, or biological sequences), by passing information from previous items in a sequence to the current item in a sequence^29^. Long Short Term Memory networks (LSTMs)^30^ are an extension of RNNs, which better learn long-term dependencies by handling the RNN tendency to “forget” information farther away in a sequence^31^. LSTMs maintain a “cell state” which contains the “memory” of the information in the previous items in the sequence. LSTMs learn additional parameters which decide at each step in the sequence which information in the “cell state” to “forget” or “update”.

LookingGlass uses a three-layer LSTM encoder model with 1,152 units in each hidden layer and an embedding size of 104 based on the results of hyperparameter tuning (see below). It divides the sequence into characters using a kmer size of 1 and a stride of 1, i.e. is a character-level language model. LookingGlass is trained in a self-supervised manner to predict a masked nucleotide, given the context of the preceding nucleotides in the sequence. For each read in the training sequence, multiple training inputs are considered, shifting the nucleotide that is masked along the length of the sequence from the second position to the final position in the sequence. Because it is a character-level model, a linear decoder predicts the next nucleotide in the sequence from the possible vocabulary items ‘A’, ‘C’, ‘G’, and ‘T’, with special tokens for ‘beginning of read’, ‘unknown nucleotide’ (for the case of ambiguous sequences), ‘end of read’ (only ‘beginning of read’ was tokenized during LookingGlass training), and a ‘padding’ token (used for classification only).

Regularization and optimization of LSTMs require special approaches to dropout and gradient descent for best performance^32^. The *fastai* library^33^ offers default implementations of these approaches for natural language text, and so we adopt the fastai library for all training presented in this paper. We provide the open-source *fastBio* python package^2^ which extends the fastai library for use with biological sequences.

LookingGlass was trained on a Pascal P100 GPU with 16GB memory on Microsoft Azure, using a batch size of 512, a back propagation through time (bptt) window of 100 base pairs, the Adam optimizer^34^, and utilizing a Cross Entropy loss function (SI Table 2). Dropout was applied at variable rates across the model (SI Table 2). LookingGlass was trained for a total of 12 days for 75 epochs, with progressively decreasing learning rates based on the results of hyperparameter optimization (see below): for 15 epochs at a learning rate of 1e-2, for 15 epochs at a learning rate of 2e-3, and for 45 epochs at a learning rate of 1e-3.

#### Hyperparameter optimization

Hyperparameters used for the final training of LookingGlass were tuned using a randomized search of hyperparameter settings. The tuned hyperparameters included kmer size, stride, number of LSTM layers, number of hidden nodes per layer, dropout rate, weight decay, momentum, embedding size, bptt size, learning rate, and batch size. An abbreviated dataset consisting of ten randomly selected read-length chunks from the *GTDB representative set* was created for testing many parameter settings rapidly. A language model was trained for two epochs for each randomly selected hyperparameter combination, and those conditions with the maximum performance were accepted. The hyperparameter combinations tested and the selected settings are described in the associated Github repository^3^.

### II. LookingGlass validation and analysis of embeddings

#### Functional relevance

##### Dataset generation

In order to assess the ability of the LookingGlass embeddings to inform the molecular function of sequences, metagenomic sequences from a diverse set of environments were downloaded from the Sequence Read Archive (SRA)^35^. We used MetaSeek^28^ to choose ten metagenomes at random from each of the ‘environmental packages’ defined by the MIxS metadata standards^36^: ‘built environment’, ‘host-associated’, ‘human-gut’, ‘microbial mat/biofilm’, ‘miscellaneous’, ‘plant-associated’, ‘sediment’, ‘soil’, ‘wastewater/sludge’, and ‘water’, for a total of 100 metagenomes. The SRA IDs used are available in (SI Table 3). The raw DNA reads for these 100 metagenomes were downloaded from the SRA with the NCBI e-utilities. These 100 metagenomes were annotated with the mi-faser tool^37^ with the --read-map option to generate predicted functional annotation labels (to the fourth digit of the Enzyme Commission (EC) number), out of 1,247 possible EC labels, for each annotatable read in each metagenome. These reads were then split 80%/20% into ‘training’/’validation candidate’ sets of reads. To ensure that there was minimal overlap in sequence similarity between the training and validation set, we compared the ‘validation candidate’ sets of each EC annotation to the training set for that EC number with CD-HIT^38^, and filtered out any reads with >80% DNA sequence similarity to the reads of that EC number in the training set (the minimum CD-HIT DNA sequence similarity cutoff). In order to balance EC classes in the training set, overrepresented ECs in the training set were downsampled to the mean count of read annotations (52,353 reads) before filtering with CD-HIT. After CD-HIT processing, any underrepresented EC numbers in the training set were oversampled to the mean count of read annotations (52,353 reads). The validation set was left unbalanced to retain a distribution more realistic to environmental settings. The final training set contained 61,378,672 reads, while the validation set contained 2,706,869 reads. We term this set of reads and their annotations the *mi-faser functional set* (Table 1).

As an external test set, we used a smaller number of DNA sequences from genes with experimentally validated molecular functions. We linked the manually curated entries of Bacterial or Archaeal proteins from the Swiss-Prot database^39^ corresponding to the 1,247 EC labels in the *mi-faser functional set* with their corresponding genes in the EMBL database^40^. We downloaded the DNA sequences, and selected ten read-length chunks at random per coding sequence. This resulted in 1,414,342 read-length sequences in the test set. We term this set of reads and their annotations the *Swiss-Prot functional set* (Table 1).

##### Fine-tuning procedure

We fine-tuned the LookingGlass language model to predict the functional annotation of DNA reads, to demonstrate the speed with which an accurate model can be trained using our pretrained LookingGlass language model. The architecture of the model retained the 3-layer LSTM encoder and the weights of the LookingGlass language model encoder, but replaced the language model decoder with a new multi-class classification layer with pooling (with randomly initialized weights). This pooling classification layer is a sequential model consisting of the following layers: a layer concatenating the output of the LookingGlass encoder with min, max, and average pooling of the outputs (for a total dimension of 104*3 = 312), a batch normalization^41^ layer with dropout, a linear layer taking the 312-dimensional output of the batch norm layer and producing a 50-dimensional output, another batch normalization layer with dropout, and finally a linear classification layer that outputs the predicted functional annotation of a read as a probability distribution of the 1,247 possible mi-faser EC annotation labels. We then trained the functional classifier on the *mi-faser functional set* described above. Because the >61 million reads in the training set were too many to fit into memory, training was done in 13 chunks of ~5-million reads each until one total epoch was completed. Hyperparameter settings for the functional classifier training are seen in SI Table 2.

##### Encoder embeddings and MANOVA test

To test whether the LookingGlass language model embeddings (before fine-tuning, above) are distinct across functional annotations, a random subset of ten reads per functional annotation was selected from each of the 100 SRA metagenomes (or the maximum number of reads present in that metagenome for that annotation, whichever was greater). This also ensured that reads were evenly distributed across environments. The corresponding fixed-length embedding vectors for each read was produced by saving the output from the LookingGlass encoder (before the embedding vector is passed to the language model decoder) for the final nucleotide in the sequence. This vector represents a contextually relevant embedding for the overall sequence. The statistical significance of the difference between embedding vectors across all functional annotation groups was tested with a MANOVA test using the R stats package^42^.

#### Evolutionary relevance

##### Dataset generation

The OrthoDB database^43^ provides orthologous groups (OGs) of proteins at various levels of taxonomic distance. For instance, the OrthoDB group ‘77at2284’ corresponds to proteins belonging to ‘Glucan 1,3-alpha-glucosidase at the Sulfolobus level’, where ‘2284’ is the NCBI taxonomy ID for the genus *Sulfolobus*.

We tested whether embedding similarity of homologous sequences (sequences within the same OG) is higher than that of nonhomologous sequences (sequences from different OGs). We tested this in OGs at multiple levels of taxonomic distance – genus, family, order, class, and phylum. At each taxonomic level, ten individual taxa at that level were chosen from across the prokaryotic tree of life (e.g. for the genus level, *Acinetobacter*, *Enterococcus*, *Methanosarcina*, *Pseudomonas*, *Sulfolobus*, *Bacillus*, *Lactobacillus*, *Mycobacterium*, *Streptomyces*, and *Thermococcus* were chosen). For each taxon, 1,000 randomly selected OGs corresponding to that taxon were chosen; for each of these OGs, five randomly chosen genes within this OG were chosen.

OrthoDB cross-references OGs to UniProt^39^ IDs of the corresponding proteins. We mapped these to the corresponding EMBL coding sequence (CDS) IDs^40^ via the UniProt database API^39^; DNA sequences of these EMBL CDSs were downloaded via the EMBL database API. For each of these sequences, we generated LookingGlass embedding vectors.

##### Homologous and nonhomologous sequence pairs

To create a balanced dataset of homologous and nonhomologous sequence pairs, we compared all homologous pairs of the five sequences in an OG (total of ten homologous pairs) to an equal number of randomly-selected out-of-OG comparisons for the same sequences; i.e., each of the five OG sequences was compared to 2 other randomly-selected sequences from any other randomly-selected OG (total of ten nonhomologous pairs). We term this set of sequences, and their corresponding LookingGlass embeddings, the *OG homolog set* (Table 1).

##### Embedding and sequence similarity

For each sequence pair, the sequence and embedding similarity were determined. The embedding similarity was calculated as the cosine similarity between embedding vectors. The sequence similarity was calculated as the Smith-Waterman alignment score using the BioPython^44^ pairwise2 package, with a gap open penalty of −10 and a gap extension penalty of −1. The IDs of chosen OGs, the cosine similarities of the embedding vectors, and sequence similarities of the DNA sequences are available in the associated Github repository^3^.

#### Environmental Relevance

##### Encoder embeddings and MANOVA test

The LookingGlass embeddings and the environment of origin for each read in the *mi-faser functional set* were used to test the significance of the difference between the embedding vectors across environmental contexts. The statistical significance of this difference was evaluated with a MANOVA test using the R stats package^42^.

### III. Oxidoreductase classifier

#### Dataset generation

The manually curated, reviewed entries of the Swiss-Prot database^39^ were downloaded (June 2, 2020). Of these, 23,653 entries were oxidoreductases (EC number 1.-.-.-) of Archaeal or Bacterial origin (988 unique ECs). We mapped their UniProt IDs to both their EMBL CDS IDs and their UniRef50 IDs via the UniProt database mapper API. Uniref50 IDs identify clusters of sequences with >50% amino acid identity. This cross-reference identified 28,149 EMBL CDS IDs corresponding to prokaryotic oxidoreductases, belonging to 5,451 unique UniRef50 clusters. We split this data into training, validation, and test sets such that each UniRef50 cluster was contained in only one of the sets, i.e. there was no overlap in EMBL CDS IDs corresponding to the same UniRef50 cluster across sets. This ensures that the oxidoreductase sequences in the validation and test sets are dissimilar to those seen during training. The DNA sequences for each EMBL CDS ID were downloaded via the EMBL database API. This data generation process was repeated for a random selection of non-oxidoreductase UniRef50 clusters, which resulted in 28,149 non-oxidoreductase EMBL CDS IDs from 13,248 unique UniRef50 clusters.

~50 read-length chunks (selected from the representative read-length distribution, as above) were selected from each EMBL CDS DNA sequence, with randomly selected start positions on the gene and a 50% chance of selecting the reverse complement, such that an even number of read-length sequences with ‘oxidoreductase’ and ‘non-oxidoreductase’ labels were generated for the final dataset. This procedure produced a balanced dataset with 2,372,200 read-length sequences in the training set, 279,200 sequences in the validation set, and 141,801 sequences in the test set. We term this set of reads and their annotations the *oxidoreductase model set* (Table 1).

#### Fine-tuning procedure

Since our functional annotation classifier addresses a closer classification task to the oxidoreductase classifier than LookingGlass itself, the architecture of the oxidoreductase classifier was fine-tuned starting from the functional annotation classifier, replacing the decoder with a new pooling classification layer (as described above for the functional annotation classifier) and with a final output size of 2 to predict ‘oxidoreductase’ or ‘not oxidoreductase’. Fine tuning of the oxidoreductase classifier layers was done successively, training later layers in isolation and then progressively including earlier layers into training, using discriminative learning rates ranging from 1e-2 to 5e-4, as previously described^45^.

#### Model performance in metagenomes

16 marine metagenomes from the surface (SRF, ~5 meters) and mesopelagic (MES, 175-800 meters) from eight stations sampled as part of the TARA expedition^46^ were downloaded from the SRA^35^ (SI Table 4, SRA accession numbers ERR598981, ERR599063, ERR599115, ERR599052, ERR599020, ERR599039, ERR599076, ERR598989, ERR599048, ERR599105, ERR598964, ERR598963, ERR599125, ERR599176, ERR3589593, and ERR3589586). Metagenomes were chosen from a latitudinal gradient spanning polar, temperate, and tropical regions and ranging from - 62 to 76 degrees latitude. Mesopelagic depths from four out of the eight stations were sampled from oxygen minimum zones (OMZs, where oxygen <20 μmol/kg). Each metagenome was rarefied to twenty million randomly selected sequences. We term this set of reads the *oxidoreductase metagenome set* (Table 1, SI Table 4). Predictions of “oxidoreductase” or “not oxidoreductase” were made for these sequences with the oxidoreductase classifier. To compare model predictions to alternative functional annotation methods, reads in the *oxidoreductase metagenome set* were annotated with mi-faser^37^ with the --read-map option, and with the MG-RAST functional annotation pipeline^47^ using default settings.

### IV. Reading Frame classifier

#### Dataset generation

For each taxonomic order, the coding sequence (CDS) files of one of the genome IDs in the *GTDB representative set* were downloaded from NCBI^27^. These were split into read-length chunks as described above. Note that because each sequence is a coding sequence, the true frame of translation for each read-length chunk was known; this translation frame label of (1, 2, 3, −1, −2, or −3) was recorded for each read-length input^3^. We term this set of reads the *reading frame set* (Table 1).

#### Fine-tuning procedure

The translation frame classifier was adjusted with a pooling classification layer with an output size of six for the six possible translation frame labels. Fine tuning was performed over successive layers with discriminative learning rates ranging from 1e-3 to 5e-5 as described for the oxidoreductase classifier.

### V. Optimal temperature classifier

#### Dataset generation

The optimal growth temperature for 19,474 microorganisms was manually curated from multiple sources: BacDive^48^, DSMZ^49^, Pasteur Institute (PI), the National Institute for Environmental Studies (NIES)^50^, and a curated list from a previous work^51^. BacDive data is available through their API, which contains calls to retrieve the species list and to get all data about a specific species. For DSMZ, PI, and NIES databases we used previously published^52^ data files (for DSMZ and PI) or scripts and method (NIES) to query optimal growth temperature information (accessed July 2020). We finally cross-referenced optimal growth temperature of these organisms to their NCBI taxonomy ID^53^.

Previous studies have shown a strong correlation between enzyme optimal temperature and organism optimal growth temperature^52^. We assumed that core housekeeping enzymes, such as those involved in transcription and translation, would have the same optimal functional temperature as the organism itself. Thus, we cross-referenced the 19,474 microorganisms identified above to the UniProt IDs belonging to those taxa for the housekeeping genes: RNA polymerase (EC 2.7.7.6), RNA helicase (EC 3.6.4.13), DNA polymerase (EC 2.7.7.7), DNA primase (EC 2.7.7.101 for Bacteria, EC 2.7.7.102 for Archaea), DNA helicase (EC 3.6.4.12), DNA ligase (ECs 6.5.1.1, 6.5.1.2, 6.5.1.6, and 6.5.1.7), and topoisomerase (ECs 5.6.2.1 and 5.6.2.2). Finally, we linked these UniProt IDs to the corresponding EMBL CDS IDs, downloaded the gene sequences, and split them into read-length chunks as described above.

The optimal temperature label for each read was derived from the optimal growth temperature from its source organism; range [4-104.5] C°. The optimal temperature labels were converted to categorical labels of ‘psychrophilic’ for optimal temperatures <15 C°, ‘mesophilic’ for [20-40] C°, and ‘thermophilic’ for >50 C°. The training, validation, and test sets were split by EC number such that only sequences from EC 3.6.4.13 were in the validation set, only sequences from EC 6.5.1.2 were in the test set, and all other EC numbers were in the training set. Finally, the inputs from each label category were either downsampled or upsampled (as described above for the *mi-faser functional set*) to a balanced number of inputs for each class. This resulted in 5,971,152 inputs in the training set with ~2,000,000 reads per label; 597,136 inputs in the validation set with ~200,000 reads per label; and 296,346 inputs to the test set with ~100,000 reads per label. We term this set of reads and their annotations the *optimal temp set* (Table 1).

#### Fine-tuning procedure

The optimal temperature classifier was adjusted with a pooling classification layer with an output size of three for the three possible optimal temperature labels, as described above. Fine tuning was performed over successive layers with discriminative learning rates ranging from 5e-2 to 5e-4 as described for the oxidoreductase classifier.

### VI. Metrics

Model performance metrics for accuracy (all classifiers), precision, recall, and F1 score (binary classifiers only) are defined as below:

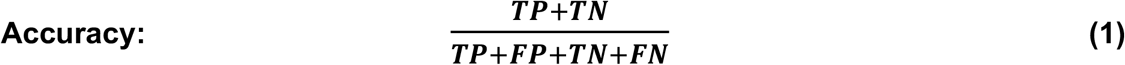

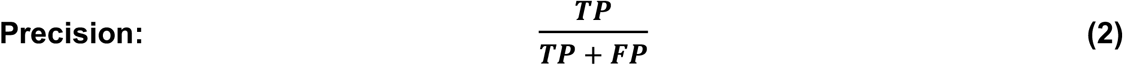

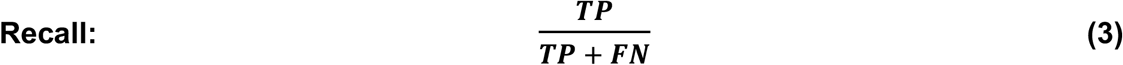

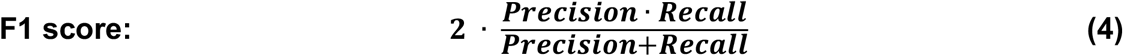

where TP is a true positive (correct positive label prediction), FP is a false positive (incorrect prediction of the positive label), TN is a true negative (correct negative label prediction), and FN is a false negative (incorrect prediction of the negative label).

### VII. Software, model deployment and reproducibility

The LookingGlass pretrained model, as well as the pretrained functional classifier, oxidoreductase classifier, optimal temperature classifier, and reading frame classifier models, are provided in the *LookingGlass* release v1.0^1^. We also provide the *fastBio* python package that extends the fastai^33^ library for custom data loading and processing functions designed for use with biological sequences^2^. The scripts used for data gathering, training of the LookingGlass model, training of models using transfer learning, and analysis of the results presented in this paper are available in the associated Github repository^3^.

## Results

### I. LookingGlass – a “universal language of life”

The LookingGlass model was trained as a 3-layer LSTM encoder chained to a decoder predicting the next (masked) nucleotide in a DNA sequence fragment, on a set of more than 6.6 million read-length sequences selected from microbial genomes spanning each taxonomic class in the microbial tree of life (Methods).

#### LookingGlass captures functionally relevant features of sequences

The LookingGlass encoder produces a fixed-length vector embedding of each sequence input. In the *mi-faser functional* validation set containing metagenomic reads with functional annotation labels (Methods), these sequence embeddings were distinct across functional annotations (MANOVA P<10^−16^) without any additional fine-tuning. Moreover, a model was fine-tuned on the *mi-faser functional set* to predict mi-faser functional annotations to the 4^th^ EC number and achieved 81.5% accuracy (Eqn 1) on the validation set in only one epoch. At coarser resolution accuracy was improved: to 83.8% at the 3^rd^ EC number (SI Fig 3); 84.4% at the 2^nd^ EC number (Fig 1b); and 87.1% at the 1^st^ EC number (Fig 1a). In testing on an experimentally-validated set of functional annotations (*Swiss-Prot functional set*; Methods), this classifier had a lower accuracy (50.8%) that was still substantially better than random (0.08%). Thus, LookingGlass captures functionally relevant features of biological sequences, (1) distinguishing between functional classes without being expressly trained to do so and (2) enabling rapid convergence on an explicit high-dimensional functional classification task at the read level.

**Figure 1:**
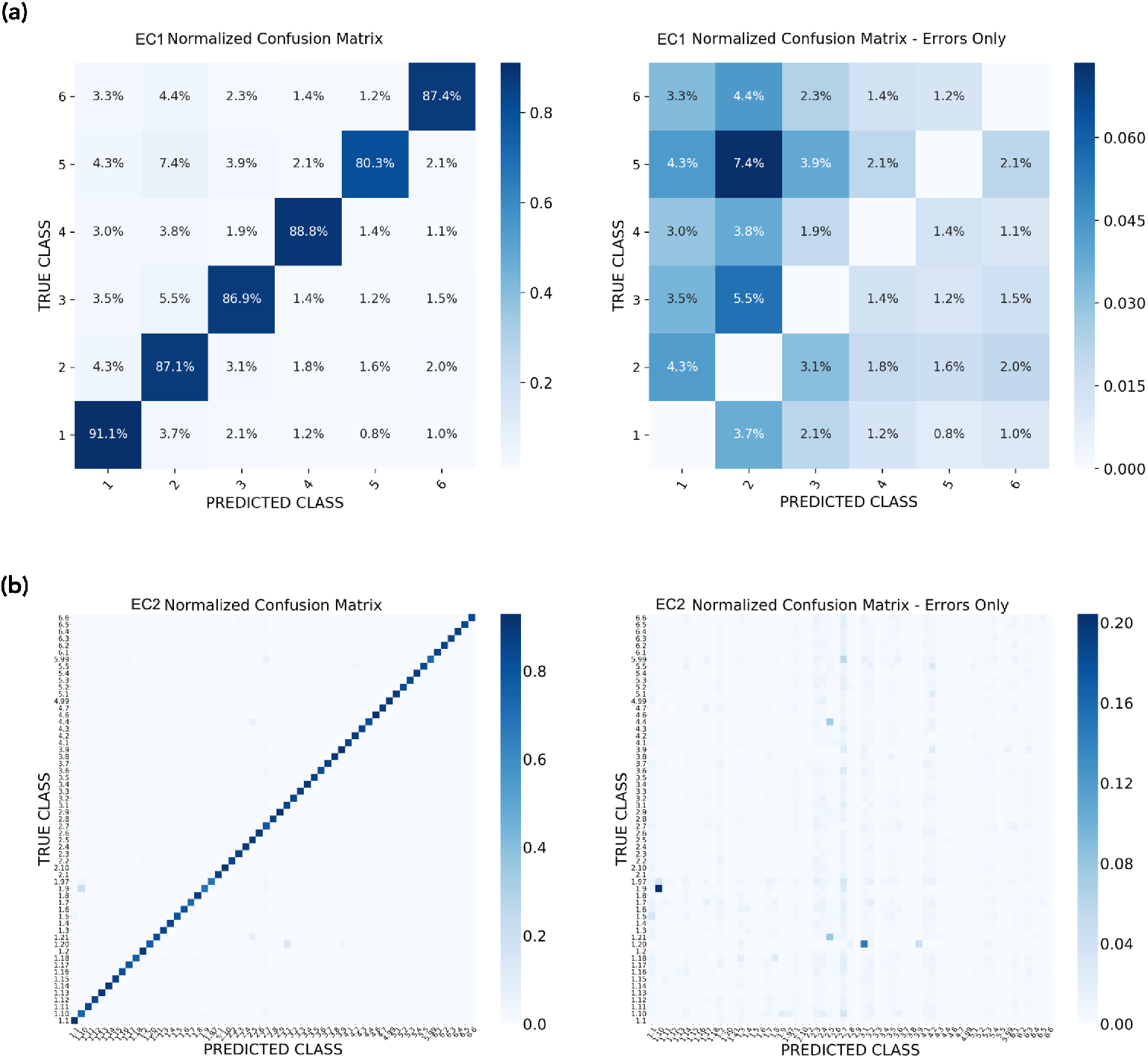
Functional annotation prediction multiclass confusion matrix. Confusion between true (y axis) and predicted (x axis) functional annotations, shown as normalized percentages of predictions for each label including correct predictions (left) and showing errors only (right), for **(a)** predictions to the 1^st^ EC number and **(b)** predictions to the 2^nd^ EC number.

#### LookingGlass captures evolutionarily-relevant features of sequences

The embedding similarity of homologous sequence pairs in the *OG homolog set* was significantly higher (unpaired t-test P<10^−16^) than that of nonhomologous pairs, with no additional fine-tuning, for fine to broad levels of phylogenetic distances, i.e. genus, family, order, class, and phylum (Fig 2a). LookingGlass embeddings differentiate homology with ~66-79% accuracy which varied by taxonomic level (SI Fig 4, SI Table 5). This variation is due to variable sequence similarity across taxa, i.e. sequences from species-level homologs have higher sequence similarity than homologs at the phylum level. Our model attained 66.4% accuracy at the phylum level (Fig 2b), 68.3% at the class level, 73.2% at the order level, 76.6% at the family level, and 78.9% at the genus level. This performance is a substantial improvement over random (50% accuracy), and was obtained from LookingGlass embeddings alone which were not expressly trained on this task.

**Figure 2:**
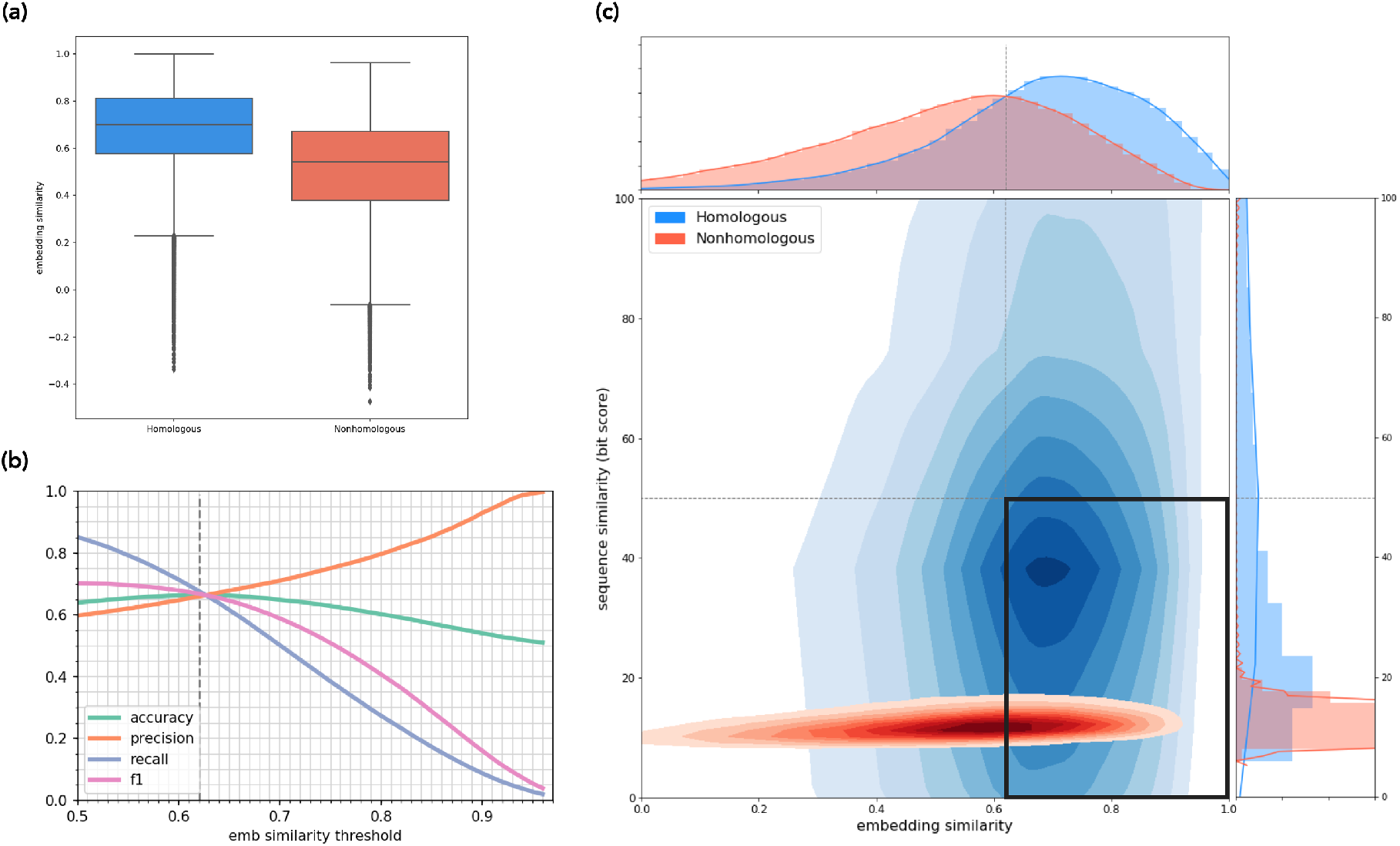
LookingGlass identifies homologous sequence pairs at the phylum level. (a) Distribution of embedding similarities for homologous (blue) and nonhomologous (red) sequence pairs are significantly different (P < 10-16). (b) Accuracy, precision, recall, and F1 metrics (Eqns 1–4) for homologous/ nonhomologous predictions across embedding similarity thresholds. Default threshold of maximum accuracy (0.62) shown in vertical dashed line. (c) Distribution of embedding and sequencing similarities for homologous (blue) and nonhomologous (red) sequence pairs. 44% of homologous sequence pairs have sequence similarity alignment scores below the threshold of 50 (horizontal line). Embedding similarity threshold (0.62, vertical line) separates homologous and nonhomologous sequence pairs with maximum accuracy. Bold black box in the lower right indicates homologous sequences correctly identified by LookingGlass that are missed using alignments.

LookingGlass embeddings differentiate between homologous and nonhomologous sequences independent of their sequence similarity (Smith-Waterman alignments, Methods). This is particularly useful since many (e.g. 44% at the phylum level, SI Table 5) homologs have very low sequence similarity (alignment score < 50; Fig 2c, SI Table 5). For these, LookingGlass embedding similarity is still high, indicating that our model captures evolutionary relationships between sequences, even where traditional algorithmic approaches do not. In fact, embedding similarity between sequences is poorly correlated with the sequence similarity alignment score (Pearson R^2^=0.28-0.44). The high accuracy with which LookingGlass identifies homologs indicates that it captures high-level features reflecting evolutionary relationships between sequences.

#### LookingGlass differentiates sequences from disparate environmental contexts

The sequences in the *mi-faser functional set* have distinct embedding fingerprints across different environments – embedding similarity between environments is generally lower than embedding similarity within an environment (Fig 3, MANOVA P<10^−16^), even though the LookingGlass embeddings were not explicitly trained to recognize environmental labels. While there is some overlap of embeddings across environmental contexts, those with the most overlap are between similar environments – for example, the colocalization of ‘wastewater/sludge’ with ‘human-gut’ and ‘built environment’ (Fig. 3b).

**Figure 3:**
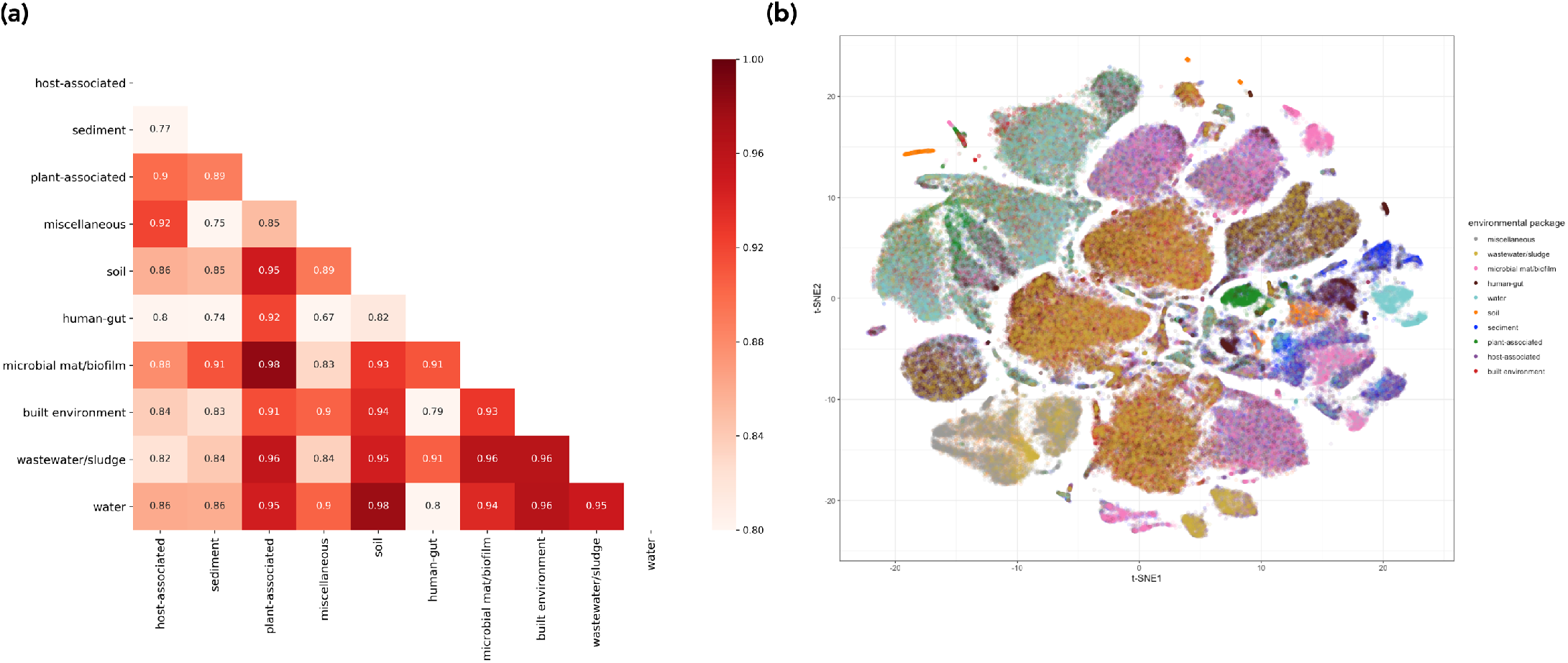
Distributions of LookingGlass embeddings across environmental packages. **(a)** Pairwise cosine similarity among the average embeddings of 20,000 randomly selected sequences from each environmental package. **(b)** t-SNE visualization of the embedding space for 20,000 randomly selected sequences from each of ten distinct environmental contexts in the *mi-faser functional* validation set. Sequences from the same environmental context generally cluster together. Colors indicate environmental package. Embeddings are significantly differentiated by environmental package (P < 10^−16^).

### II. LookingGlass enables diverse downstream transfer learning tasks

#### Mining environmental settings for functional descriptions of “microbial dark matter”

##### Using LookingGlass and transfer learning to identify novel functional groups

By using LookingGlass as a starting point, we can converge more quickly and with less data on a more accurate model for assigning molecular functions at the read level. Additionally, downstream models addressing similar tasks can in turn be used as pretrained models for further fine-tuning. To demonstrate this, we fine-tuned the LookingGlass functional classifier (described above) to predict whether a read-length DNA sequence likely comes from an oxidoreductase-encoding gene (EC number 1.-.-.-). Our fine-tuned model was able to correctly classify previously unseen (<50% amino acid sequence-identical) oxidoreductases with 82.3% accuracy at the default prediction threshold of 0.5 (Fig 4). Oxidoreductases are a deeply branched, highly diverse class of enzymes, such that sequence similarity within a single functional annotation (EC number) is often very low; the DNA sequence identity of oxidoreductase gene sequences within a single EC number in the *oxidoreductase model* validation set was a median of 59%, and was as low as 17%. As such, oxidoreductases can be difficult to identify via sequence similarity-based homology searches in environmental samples (e.g. box in Fig 2c). The oxidoreductase classifier, in contrast, achieves high model performance even in such cases where sequence similarity within EC annotations is low. Notably, the average model performance for a given EC number was independent of the sequence similarity of genes within that EC (R^2^=0.004, SI Fig 5).

**Figure 4:**
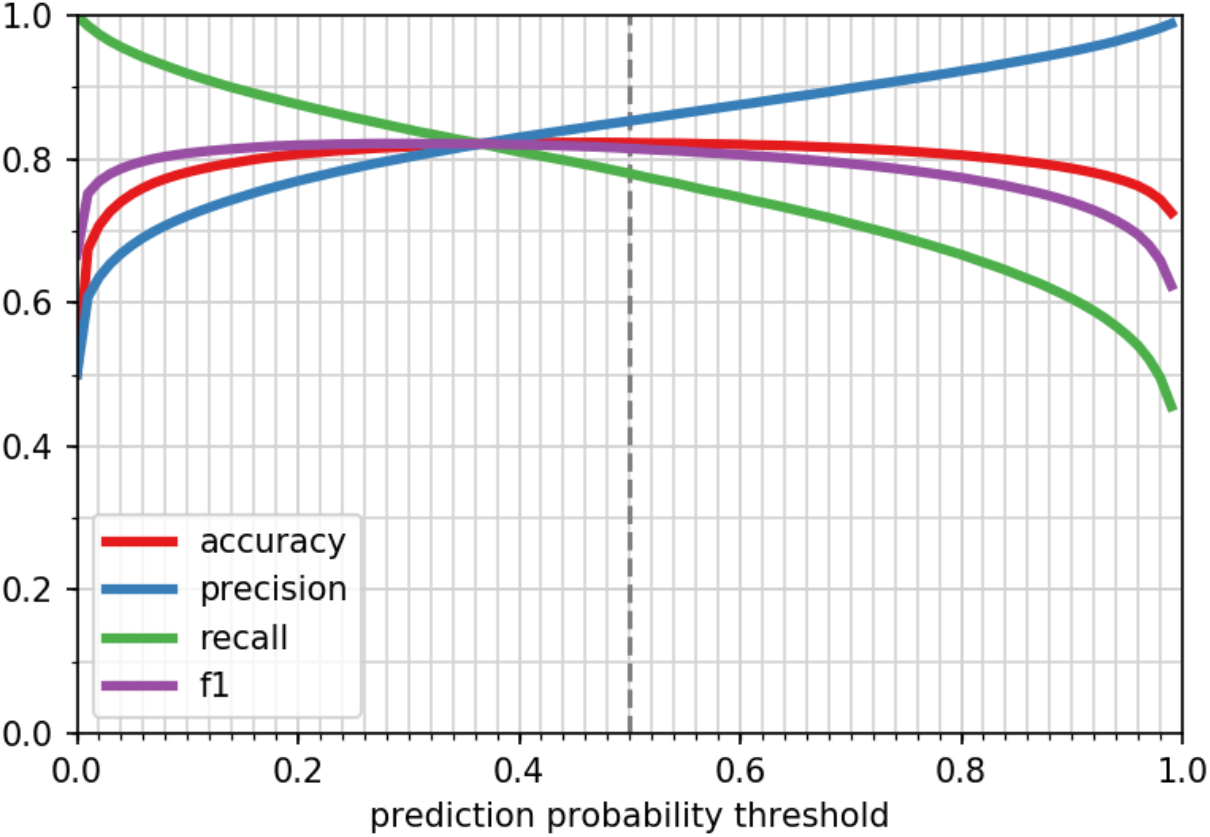
Performance of the oxidoreductase classifier. Accuracy, precision, recall, and F1 score metrics (Eqns 1–4) of the oxidoreductase classifier across prediction probability thresholds. Default threshold of 0.5 shown in vertical dashed line

##### Mining novel, unannotated oxidoreductases from metagenomes along a latitudinal and depth gradient in the global ocean

The majority of sequencing reads from environmental metagenomes are routinely unable to be functionally annotated^54^. To demonstrate the advantage of the oxidoreductase classifier over traditional homology-based approaches, we evaluated our model on twenty million randomly-selected reads from each of 16 marine metagenomes in the *oxidoreductase metagenome* set spanning broad ranges in latitude (from −62 to 76 degrees), depth (from the surface, ~5 meters, to mesopelagic, ~200-1,000 meters), and oxygen concentrations (including four mesopelagic samples from oxygen minimum zones).

The percentage of reads predicted to be oxidoreductases ranged from 16.4% - 20.6%, and followed trends with depth and latitude (Fig 5). The relative abundance of oxidoreductases was significantly higher in mesopelagic depths than in surface waters (Fig 5a, ANOVA P=0.02), with marginally higher (albeit not statistically significant) proportions of oxidoreductases in the oxygen minimum zones relative to oxygen-replete mesopelagic samples (P=0.13). There was also a significant trend in the relative abundance of oxidoreductases along latitudinal gradients in surface waters (Fig 5b, R^2^=0.79, P=0.04), with higher proportions of oxidoreductases in higher latitudes. This latitudinal trend was reflected in a similar, but weaker, temperature-driven trend (R^2^= −0.66, P=0.11, SI Fig 6).

**Figure 5:**
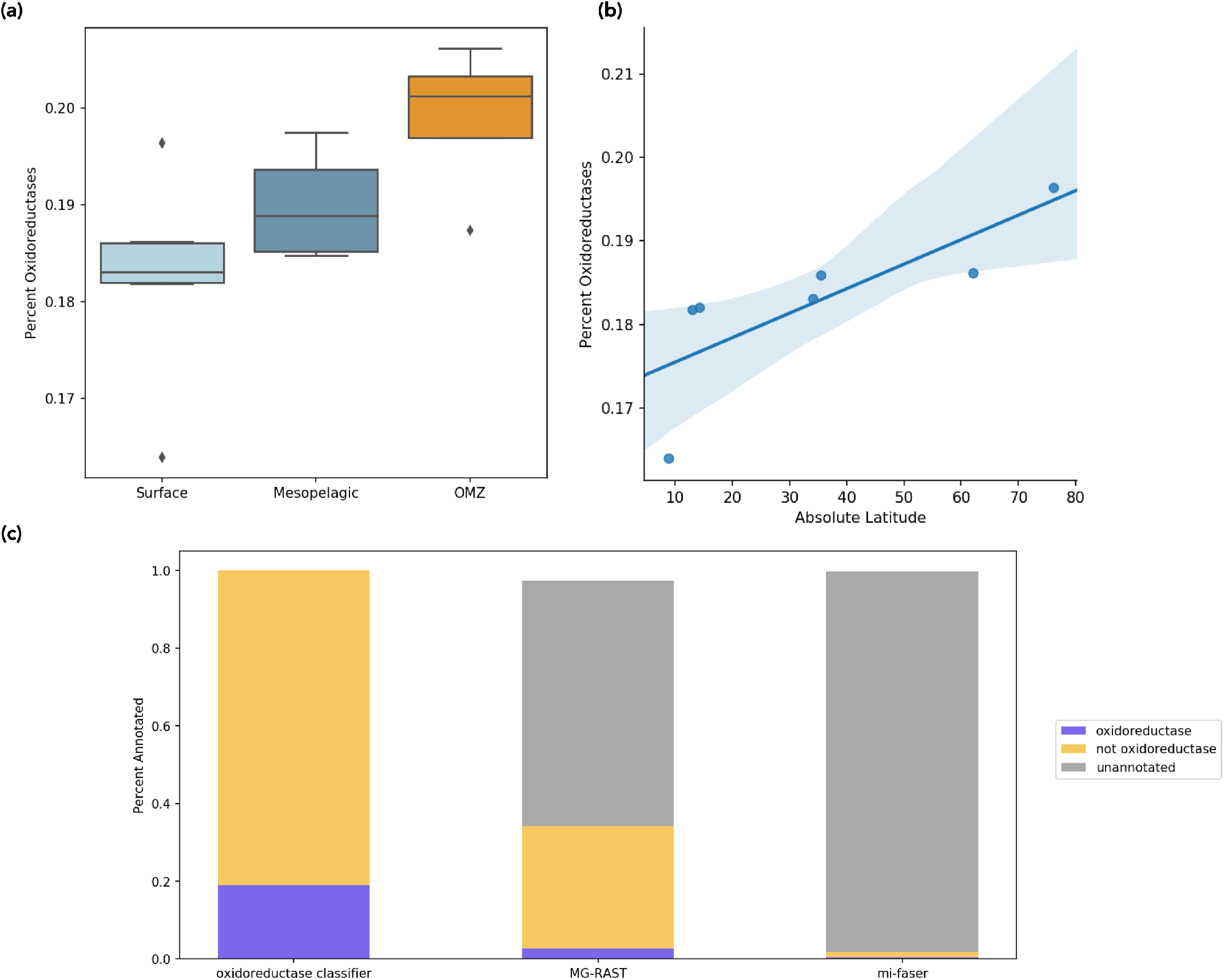
Oxidoreductase identification in marine metagenomes. **(a)** Proportion of oxidoreductase sequences (y axis) predicted by the oxidoreductase classifier in surface, mesopelagic, and oxygen minimum zone (OMZ) depths. **(b)** Correlation between the proportion of oxidoreductases and absolute degrees latitude in surface metagenomes of the *oxidoreductase metagenome set* (R^2^=0.79, P=0.04). **(c)** Proportion of sequences predicted as oxidoreductases, not oxidoreductases, or left unannotated across the oxidoreductase classifier, MG-RAST, and mi-faser tools.

Two alternative functional annotation tools, mi-faser^37^ and MG-RAST^47^, were only able to annotate a much smaller proportion of sequences in these metagenomes (Fig 5c, SI Table 6), with even smaller proportions of oxidoreductases identified. MG-RAST annotated 26.7-50.3% of the reads across metagenomes, with 0.01-4.0% of reads identified as oxidoreductases. Mi-faser annotated 0.17-2.9% of the reads, of which 0.04-0.59% were oxidoreductases. In both cases, a majority of reads remained unannotated, a condition typical of homology-based functional annotation approaches^54^. As a result, a large proportion of enzymes in the environment are unlikely to be recovered using these approaches, which may also skew the observed trends across samples. Notably, the depth and latitudinal trends identified with the oxidoreductase classifier were not reported by either MG-RAST or mi-faser (SI Fig 7). There was no significant difference in the proportion of oxidoreductases predicted in the surface vs. mesopelagic waters for either MG-RAST (P=0.73) or mi-faser (P=0.60) and no significant correlation with latitude in surface waters for either mi-faser (R^2^=0.58, P=0.17) or MG-RAST (R^2^= −0.49, P=0.27); note that MG-RAST in fact observed an anticorrelation trend for the latter (although still insignificant). This highlights the potential importance of unannotatable reads in driving functional patterns in the environment, which can be captured by the approach and models described here and would otherwise be missed using traditional approaches.

#### Reference-free translation of read-length DNA sequences to peptides

While the amino acid sequence encoded in short DNA reads is difficult to infer directly using traditional bioinformatic approaches, it is also a product of the non-random organization of DNA sequences. We fine-tuned the LookingGlass encoder to predict the translation frame start position (1, 2, 3, −1, −2, or −3) directly from read-length DNA coding sequences. This reading frame classifier attained 97.8% accuracy, a major improvement over random (16.7% accuracy). Note this classifier was trained only on coding sequences and is currently intended only for prokaryotic sources with low amounts of noncoding DNA^55^.

#### Prediction of enzyme optimal temperature from DNA sequence fragments

The optimal temperature of an enzyme is in part dependent on DNA sequence features^56,57^, but is difficult to predict, particularly from short reads. We fine-tuned LookingGlass to predict whether a read-length DNA sequence originates from an enzyme with an optimal temperature that is psychrophilic (<15 C°), mesophilic (20-40 C°), or thermophilic (>50 C°). The optimal temperature classifier was able to predict the optimal temperature category correctly with 70.1% accuracy (random accuracy =33.3%).

## Discussion

Microbes perform a vast diversity of functional roles in natural environments as well as in industrial and biomedical settings. They play a central role in regulating Earth’s biogeochemical cycles^58^, and have a tremendous impact on the health of their human hosts^59^, but the complex functional networks that drive their activities are poorly understood. Microbial genomes record a partial history of the evolution of life on Earth^60^, but much of this information is inadequately captured by homology-based inference. Microbial communities are a subject of great interest for developing natural^61^ and synthetic^62^ products for bioengineering applications, but our ability to describe, model, and manipulate the systems-level functions of these microbiomes is limited.

The LookingGlass ‘universal language of life’ creates representations of DNA sequences that capture their functional and evolutionary relevance, independent of whether the sequence is contained in reference databases. The vast majority of microbial diversity is uncultured and unannotated^4–6^. LookingGlass opens the door to harnessing the potential of this “microbial dark matter” to improve our understanding of, and ability to manipulate, microbial systems. It is a broadly useful, ‘universal’ model for downstream transfer learning tasks, enabling a wide diversity of functional predictions relevant to environmental metagenomics, bioengineering, and biomedical applications.

We demonstrate here the ability of LookingGlass to be fine-tuned to identify novel oxidoreductases, even those with low sequence similarity to currently known oxidoreductases. Applying the oxidoreductase classifier to 16 marine metagenomes identified patterns in the relative abundance of oxidoreductases that follow global gradients in latitude and depth. These observations are in line with previous studies that have identified greater overall functional and taxonomic richness^46,63^, as well as a greater diversity of oxidoreductases specifically^64^, in deep marine waters relative to shallow depths. Studies conflict, however, about whether taxonomic and functional diversity increases^63,65–67^ or decreases^68–70^ with absolute latitude. Notably, neither the latitudinal nor depth trends in oxidoreductase relative abundance observed by the oxidoreductase classifier were captured by traditional homology-based functional annotation tools. The inconsistent results produced by traditional annotation tools in this study and others further demonstrates the importance of unannotated functional diversity for cross-sample comparisons, and the potential of the approach described in this study.

There may be multiple ecological mechanisms driving the observed latitudinal and depth patterns in oxidoreductase relative abundance; for example, the streamlining of genomes^71^ that preserves oxidoreductases relative to less essential genes under resource limitation or temperature stress, or a reflection of a higher abundance of anaerobic respiration genes in mesopelagic waters relative to surface waters^72^. Future efforts to capture and compare the full functional diversity of environmental settings using the approaches described here can further illuminate and differentiate between these mechanisms.

The reads predicted to be from novel oxidoreductases are candidates for further functional characterization, and for targeted assembly of novel oxidoreductase genes. Shining light on these “dark matter” oxidoreductases can enable more complete comparisons of oxidoreductase composition and diversity across environmental gradients. Future efforts to fine tune LookingGlass for additional functional targets can expand the classes of enzymes identified and create a fuller picture of microbial functional diversity in environmental settings. By definition, poorly-studied environments contain the greatest amount of unknown functional diversity, and a tool such as LookingGlass provides a novel and important way to evaluate this functional diversity.

LookingGlass was also fine-tuned to correctly identify the reading frame, and thus the amino acid translation, of short-read DNA coding sequences. Translated amino acid sequences are used for a variety of bioinformatics applications, most notably for molecular function annotation. There are two categories of function annotation tools – those that annotate from short sequencing reads directly^37,47,73,74^ and those that annotate from assembled genes/contigs^47,75^. In both cases, DNA reads must first be converted to amino acid sequences. For short-read annotation tools, six-frame translation of each DNA sequence produces all six possible amino acid sequences for alignment to reference databases, which increases the computational burden of alignment six-fold. For tools that annotate from assemblies, datasets are first assembled and open reading frames (ORFs) predicted before amino acid sequences can be inferred. This procedure is computationally intensive, error-prone, and throws away reads that can’t be assembled or for which coding regions can’t be identified, particularly for members of the rare biosphere or in highly diverse environments. Direct translation from DNA reads thus could enable much more efficient computation for any bioinformatics application that uses read-derived amino acid sequences. Note that the reading frame classifier described here focuses on prokaryotic genomes, which generally have only ~12-14% noncoding DNA^55^. For eukaryotes, a classifier will need to be created to distinguish between coding and noncoding DNA and predict reading frames for only the coding sequences.

Finally, we demonstrated the ability of LookingGlass to be fine tuned to predict optimal enzyme temperatures from DNA sequences. Importantly, this was possible from short reads alone, although a classifier trained on assembled genes would likely yield even better results. This result demonstrates that LookingGlass can be used to discover environmentally relevant features, as well as evolutionary and functional ones. Our optimal temperature classifier may be useful across both academic and commercial applications – for instance, to compare the optimal temperatures of microbial communities across environmental gradients in temperature or geochemistry, or to identify candidate proteins of a particular function and optimal temperature of interest for industrial applications. In addition, it may also be possible to adapt the optimal temperature classifier presented here as a generative model to guide protein design of a desired function and optimal temperature.

The LookingGlass model, and the framework for transfer learning presented here, provides a framework for future efforts toward modelling of complex biological systems. LookingGlass captures the complexity of biology and its interactions with the environment, leveraging the full potential of the functional information contained in the massive amount of sequencing data being generated by the scientific community. The LookingGlass model presented here focuses on Bacterial and Archaeal DNA sequences, but low hanging fruit may include a specialized Eukaryotic DNA model, a model specific to the human genome, or a model specialized to a particular environment such as the human gut or soil microbiome. As the scientific community continues to grapple with new approaches to represent and model biological systems in ways that harness the full potential of our expanding data resources, we hope that LookingGlass can provide a foundation for transfer learning-based exploration of life on Earth.

## Acknowledgements

The authors would like to thank Paul Falkowski and the rest of the Rutgers ENIGMA team for productive discussions of the deep transfer learning approach and inspiration for downstream applications of the LookingGlass model.

## Author Contributions

AH conceived of the project, compiled data, carried out training, validation, and application of models, and deployed open source code and software. YB provided feedback throughout the project. AA and GF curated the optimal growth temperature dataset. All authors contributed to writing of the manuscript.

## References

1. Hoarfrost, A. LookingGlass release v1.0. https://github.com/ahoarfrost/LookingGlass/. (2020). doi:10.5281/zenodo.4382930

2. Hoarfrost, A. fastBio: deep learning for biological sequences. Github repository and python package. https://github.com/ahoarfrost/fastBio/. (2020). doi:10.5281/zenodo.4383283

3. Hoarfrost, A. Github repository - LoL: learning the Language of Life. https://github.com/ahoarfrost/LoL/. doi:10.5281/zenodo.4362588

4. Lloyd, K. G., Steen, A. D., Ladau, J., Yin, J. & Crosby, L. Phylogenetically Novel Uncultured Microbial Cells Dominate Earth Microbiomes. mSystems 3, e00055–18 (2018).

5. Steen, A. D. et al. High proportions of bacteria and archaea across most biomes remain uncultured. ISME J. 13, 3126–3130 (2019).

6. Lobb, B., Tremblay, B. J. M., Moreno-Hagelsieb, G. & Doxey, A. C. An assessment of genome annotation coverage across the bacterial tree of life. Microb. Genomics 6, (2020).

7. Metagenomics versus Moore’s law. Nat. Methods 6, 623 (2009).

8. Eraslan, G., Avsec, Ž., Gagneur, J. & Theis, F. J. Deep learning: new computational modelling techniques for genomics. Nat. Rev. Genet. 20, 389–403 (2019).

9. Thrun, S. Is Learning The n-th Thing Any Easier Than Learning The First? Adv. Neural Inf. Process. Syst. 7 (1996). doi:10.1.1.44.2898

10. Pan, S. J. & Yang, Q. A survey on transfer learning. IEEE Trans. Knowl. Data Eng. 22, 1345–1359 (2010).

11. Yosinski, J., Clune, J., Bengio, Y. & Lipson, H. How transferable are features in deep neural networks? Adv. Neural Inf. Process. Syst. 1–9 (2014).

12. Devlin, J., Chang, M. W., Lee, K. & Toutanova, K. BERT: Pre-training of deep bidirectional transformers for language understanding. NAACL HLT 2019 - 2019 Conf. North Am. Chapter Assoc. Comput. Linguist. Hum. Lang. Technol. - Proc. Conf. 1, 4171–4186 (2019).

13. Liu, H., Perl, Y. & Geller, J. Transfer Learning from BERT to Support Insertion of New Concepts into SNOMED CT. AMIA … Annu. Symp. proceedings. AMIA Symp. 2019, 1129–1138 (2019).

14. Peng, Y., Yan, S. & Lu, Z. Transfer Learning in Biomedical Natural Language Processing: An Evaluation of BERT and ELMo on Ten Benchmarking Datasets. 58–65 (2019). doi:10.18653/v1/w19-5006

15. Fofanov, Y. et al. How independent are the appearances of n-mers in different genomes? Bioinformatics 20, 2421–2428 (2004).

16. Kelley, D. R., Snoek, J. & Rinn, J. L. Basset: learning the regulatory code of the accessible genome with deep convolutional neural networks. 990–999 (2016). doi:10.1101/gr.200535.115.Freely

17. Taroni, J. N. et al. MultiPLIER: A Transfer Learning Framework for Transcriptomics Reveals Systemic Features of Rare Article MultiPLIER: A Transfer Learning Framework for Transcriptomics Reveals Systemic Features of Rare Disease. Cell Syst. 8, 380–394.e4 (2019).

18. Menegaux, R. & Vert, J. P. Continuous Embeddings of DNA Sequencing Reads and Application to Metagenomics. J. Comput. Biol. 26, 509–518 (2019).

19. ElAbd, H. et al. Amino acid encoding for deep learning applications. BMC Bioinformatics 21, 235 (2020).

20. Viehweger, A., Krautwurst, S., Parks, D. H., König, B. & Marz, M. An encoding of genome content for machine learning. bioRxiv 524280 (2019). doi:10.1101/524280

21. Heinzinger, M. et al. Modeling aspects of the language of life through transfer-learning protein sequences. BMC Bioinformatics 20, 1–17 (2019).

22. Alley, E. C., Khimulya, G., Biswas, S., AlQuraishi, M. & Church, G. M. Unified rational protein engineering with sequence-based deep representation learning. Nat. Methods 16, 1315–1322 (2019).

23. Rao, R. et al. Evaluating Protein Transfer Learning with TAPE. 33rd Conf. Neural Inf. Process. Syst. (NeurIPS 2019) (2019). doi:10.1101/676825

24. Rives, A. et al. Biological Structure and Function Emerge from Scaling Unsupervised Learning to 250 Million Protein Sequences. bioRxiv 1–31 (2019). doi:https://doi.org/10.1101/622803

25. Bennett, S. Solexa Ltd. Pharmacogenomics 5, 433–8 (2004).

26. Parks, D. H. et al. A standardized bacterial taxonomy based on genome phylogeny substantially revises the tree of life. Nat. Biotechnol. 36, 996 (2018).

27. Agarwala, R. et al. Database resources of the National Center for Biotechnology Information. Nucleic Acids Res. 46, D8–D13 (2018).

28. Hoarfrost, A., Brown, N., Brown, C. T. & Arnosti, C. Sequencing data discovery with MetaSeek. Bioinformatics 35, 4857–4859 (2019).

29. Jordan, M. I. Attractor dynamics and parallelism in a connectionist sequential machine. Proc. Cogn. Sci. Soc. 531–546 (1986).

30. Hochreiter, S. & Schmidhuber, J. Long Short-Term Memory. Neural Comput. 9, 1735–1780 (1997).

31. Yoshua Bengio, Patrice Simard & Paolo Frasconi. Learning Long-term Dependencies with Gradient Descent is Difficult. IEEE Trans. Neural Netw. 5, 157 (2014).

32. Merity, S., Keskar, N. S. & Socher, R. Regularizing and Optimizing LSTM Language Models. (2015).

33. Howard, J. & Gugger, S. Fastai: A layered api for deep learning. arXiv (2020). doi:10.3390/info11020108

34. Kingma, D. P. & Ba, J. L. Adam: A Method for Stochastic Optimization. 1–15 (2015).

35. Leinonen, R., Sugawara, H. & Shumway, M. The sequence read archive. Nucleic Acids Res. 39, 2010–2012 (2011).

36. Yilmaz, P. et al. Minimum information about a marker gene sequence (MIMARKS) and minimum information about any (x) sequence (MIxS) specifications. Nat. Biotechnol. 29, 415–420 (2011).

37. Zhu, C. et al. Functional sequencing read annotation for high precision microbiome analysis. Nucleic Acids Res. 46, (2018).

38. Li, W. & Godzik, A. Cd-hit: A fast program for clustering and comparing large sets of protein or nucleotide sequences. Bioinformatics 22, 1658–1659 (2006).

39. Consortium, T. U. UniProt: A worldwide hub of protein knowledge. Nucleic Acids Res. 47, D506–D515 (2019).

40. Kanz, C. et al. The EMBL nucleotide sequence database. Nucleic Acids Res. 33, 29–33 (2005).

41. Ioffe, S. & Szegedy, C. Batch Normalization: Accelerating Deep Network Training by Reducing Internal Covariate Shift. (2015).

42. Team, R. C. R: A language and environment for statistical computing. (2017).

43. Kriventseva, E. V et al. OrthoDB v8: Update of the hierarchical catalog of orthologs and the underlying free software. Nucleic Acids Res. 43, D250–D256 (2015).

44. Cock, P. J. A. et al. Biopython: Freely available Python tools for computational molecular biology and bioinformatics. Bioinformatics 25, 1422–1423 (2009).

45. Howard, J. & Ruder, S. Universal Language Model Fine-tuning for Text Classification. arXiv (2018). doi:arXiv:1801.06146v3

46. Sunagawa, S. et al. Structure and function of the global ocean microbiome. Science (80-.). 348, 1–10 (2015).

47. Meyer, F. et al. The metagenomics RAST server - A public resource for the automatic phylogenetic and functional analysis of metagenomes. BMC Bioinformatics 9, 1–8 (2008).

48. Reimer, L. C. et al. BacDive in 2019: Bacterial phenotypic data for High-throughput biodiversity analysis. Nucleic Acids Res. 47, D631–D636 (2019).

49. Parte, A. C., Carbasse, J. S., Meier-Kolthoff, J. P., Reimer, L. C. & Göker, M. List of prokaryotic names with standing in nomenclature (LPSN) moves to the DSMZ. Int. J. Syst. Evol. Microbiol. 70, 5607–5612 (2020).

50. Kawachi, M. & Noël, M. H. Microbial Culture Collection at the National Institute for Environmental Studies, Tsukuba, Japan. PICES Press 22, 43 (2014).

51. Aptekmann, A. A. & Nadra, A. D. Core promoter information content correlates with optimal growth temperature. Sci. Rep. 8, 1–7 (2018).

52. Engqvist, M. K. M. Correlating enzyme annotations with a large set of microbial growth temperatures reveals metabolic adaptations to growth at diverse temperatures 06 Biological Sciences 0605 Microbiology 06 Biological Sciences 0601 Biochemistry and Cell Biology. BMC Microbiol. 18, 1–14 (2018).

53. Wheeler, D. L. et al. Database Resources of the National Center for Biotechnology Information. Nucleic Acids Res. 33, D39–D45 (2016).

54. Tamames, J., Cobo-Simón, M. & Puente-Sánchez, F. Assessing the performance of different approaches for functional and taxonomic annotation of metagenomes. BMC Genomics 20, 1–16 (2019).

55. Konstantinidis, K. T. & Tiedje, J. M. Trends between gene content and genome size in prokaryotic species with larger genomes. Proc. Natl. Acad. Sci. U. S. A. 101, 3160–3165 (2004).

56. Sheridan, P. P., Panasik, N., Coombs, J. M. & Brenchley, J. E. Approaches for deciphering the structural basis of low temperature enzyme activity. Biochim. Biophys. Acta - Protein Struct. Mol. Enzymol. 1543, 417–433 (2000).

57. Li, W. F., Zhou, X. X. & Lu, P. Structural features of thermozymes. Biotechnol. Adv. 23, 271–281 (2005).

58. Falkowski, P. G., Fenchel, T. & Delong, E. F. The microbial engines that drive Earth’s biogeochemical cycles. Science 320, 1034–9 (2008).

59. Clemente, J. C., Ursell, L. K., Parfrey, L. W. & Knight, R. The impact of the gut microbiota on human health: An integrative view. Cell 148, 1258–1270 (2012).

60. Hug, L. et al. A new view of the tree of life. Nat Microbiol 1, 16048 (2016).

61. Pham, J. V. et al. A review of the microbial production of bioactive natural products and biologics. Front. Microbiol. 10, (2019).

62. Song, H., Ding, M. Z., Jia, X. Q., Ma, Q. & Yuan, Y. J. Synthetic microbial consortia: From systematic analysis to construction and applications. Chem. Soc. Rev. 43, 6954–6981 (2014).

63. Salazar, G. et al. Gene Expression Changes and Community Turnover Differentially Shape the Global Ocean Metatranscriptome. Cell 179, 1068–1083.e21 (2019).

64. Ramírez-Flandes, S., González, B. & Ulloa, O. Redox traits characterize the organization of global microbial communities. Proc. Natl. Acad. Sci. U. S. A. 116, 3630–3635 (2019).

65. Fuhrman, J. A. et al. A latitudinal diversity gradient in planktonic marine bacteria. Proc. Natl. Acad. Sci. U. S. A. 105, 7774–8 (2008).

66. Ibarbalz, F. M. et al. Global Trends in Marine Plankton Diversity across Kingdoms of Life AR OCEANS EXPEDITION Article Global Trends in Marine Plankton Diversity across Kingdoms of Life. 1084–1097 (2019). doi:10.1016/j.cell.2019.10.008

67. Sul, W. J., Oliver, T. A., Ducklow, H. W., Amaral-Zettlera, L. A. & Sogin, M. L. Marine bacteria exhibit a bipolar distribution. Proc. Natl. Acad. Sci. U. S. A. 110, 2342–2347 (2013).

68. Ghiglione, J.-F. et al. Pole-to-pole biogeography of surface and deep marine bacterial communities. Proc. Natl. Acad. Sci. U. S. A. 109, 17633–8 (2012).

69. Ladau, J. et al. Global marine bacterial diversity peaks at high latitudes in winter. ISME J. 7, 1669–77 (2013).

70. Raes, E. J. et al. Oceanographic boundaries constrain microbial diversity gradients in the south pacific ocean. Proc. Natl. Acad. Sci. U. S. A. 115, E8266–E8275 (2018).

71. Giovannoni, S. J., Cameron Thrash, J. & Temperton, B. Implications of streamlining theory for microbial ecology. ISME J. 8, 1553–1565 (2014).

72. Ulloa, O., Canfield, D. E., DeLong, E. F., Letelier, R. M. & Stewart, F. J. Microbial oceanography of anoxic oxygen minimum zones. Proc. Natl. Acad. Sci. U. S. A. 109, 15996–16003 (2012).

73. Buchfink, B., Xie, C. & Huson, D. H. Fast and sensitive protein alignment using DIAMOND. Nat. Methods 12, 59–60 (2014).

74. Nazeen, S., Yu, Y. W. & Berger, B. Carnelian uncovers hidden functional patterns across diverse study populations from whole metagenome sequencing reads. Genome Biol. 21, 1–18 (2020).

75. Seemann, T. Prokka: Rapid prokaryotic genome annotation. Bioinformatics 30, 2068–2069 (2014).

