## Supplemental Information for "Shedding Light on Microbial Dark Matter with A Universal Language of Life"

### Supporting Online Material for the manuscript entitled: Shedding Light on Microbial Dark Matter with A Universal Language of Life

**Hoarfrost, A., Aptekmann, A., Farfañuk, G., Bromberg, Y.**

#### Table of Contents for Supporting Online Material

**Supplemental Figures 1-7 ........................................................................................ p.1-5**

**Supplemental Tables 1-6 ........................................................................................ p.6-11**

**References for Supporting Online Material ..................................................................... p.11**

#### Supplemental Figures

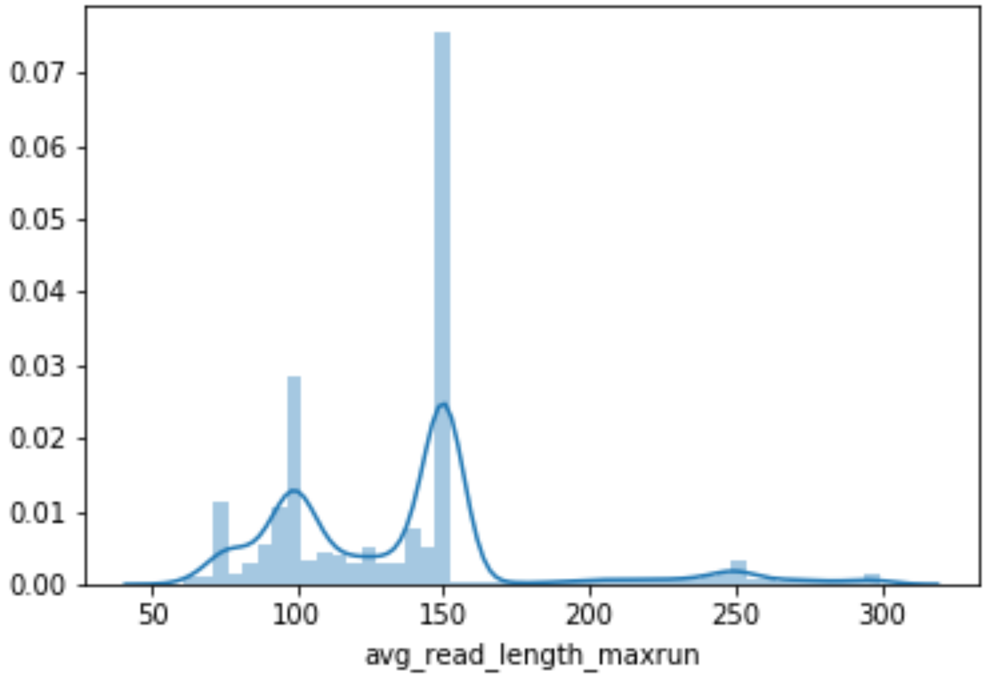


**SI Fig. 1** – The average read length distribution of the sequencing data of the 7,909 genomes with available metadata in the Genome Taxonomy Database (GTDB)[1].

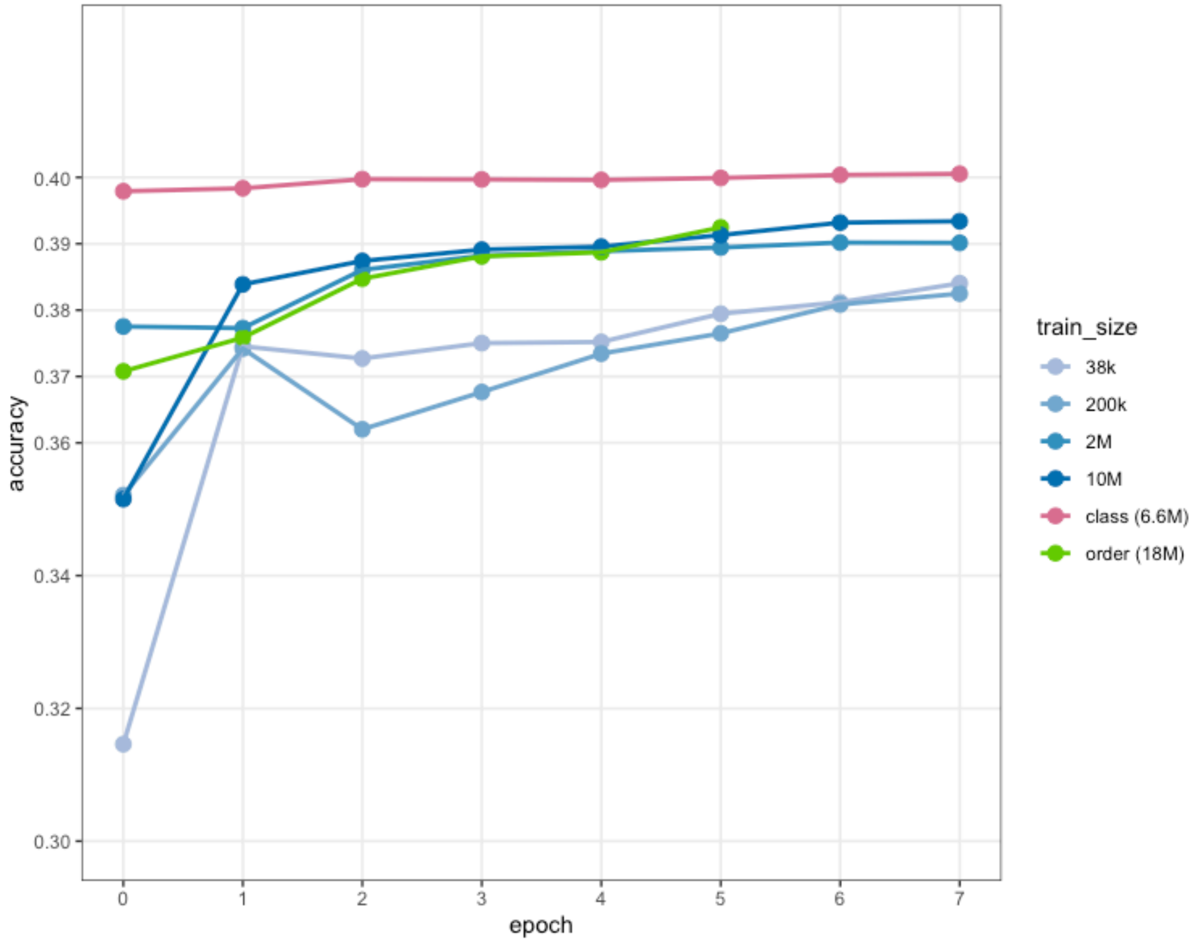


**SI Fig. 2** – Model performance over successive epochs for progressively larger datasizes randomly subsampled from the full representative GTDB genome set (blue lines), and for all reads sourced from randomly selected genomes evenly distributed across the GTDB taxonomic tree at the order level (green) and the class level (pink). Class-level partitioning of data enables much faster convergence on maximum performance.

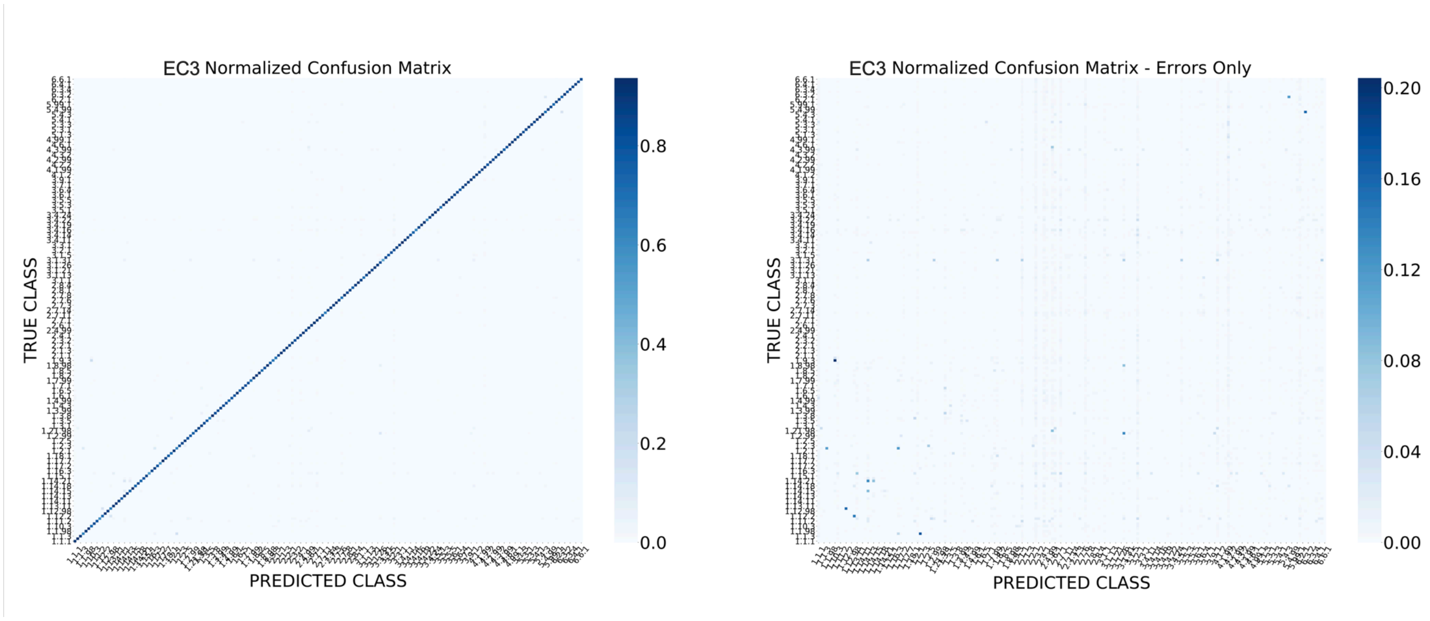


**SI Fig. 3** - Confusion between true (y axis) and predicted (x axis) functional annotations for the functional classifier, shown as normalized percentages of predictions for each label including correct predictions (left) and showing errors only (right), for predictions to the 3rd EC number.

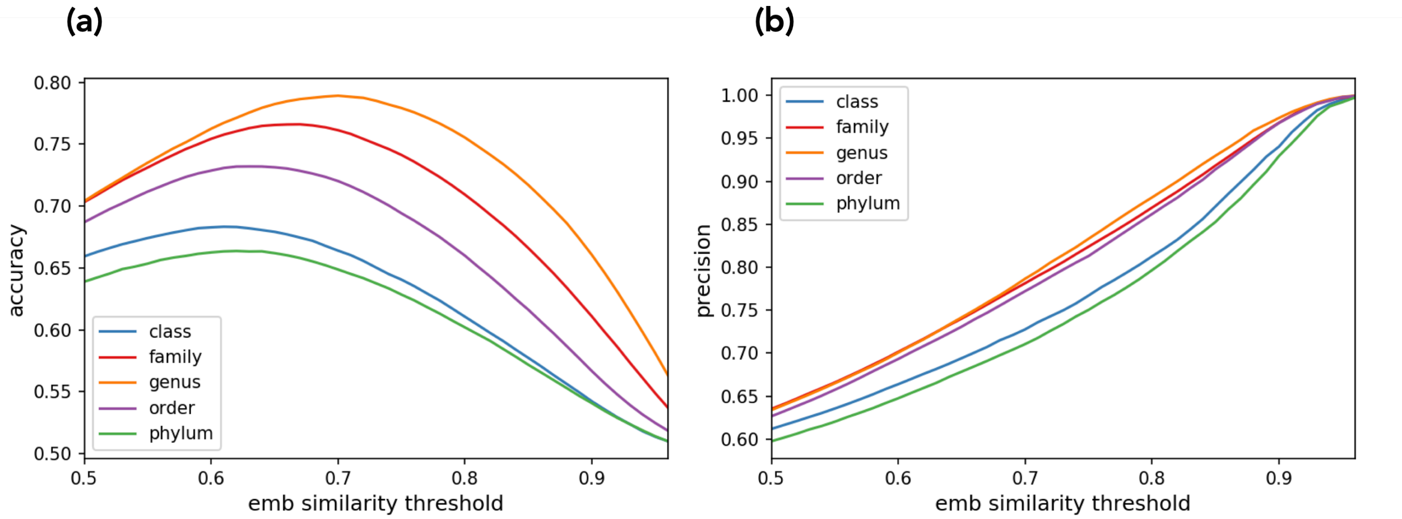


**SI Fig. 4** – (a) Accuracy and (b) precision of predictions of whether two sequences are 'homologous' or 'nonhomologous', relative to the embedding cosine similarity threshold chosen, for each of the considered levels of taxonomic specificity.

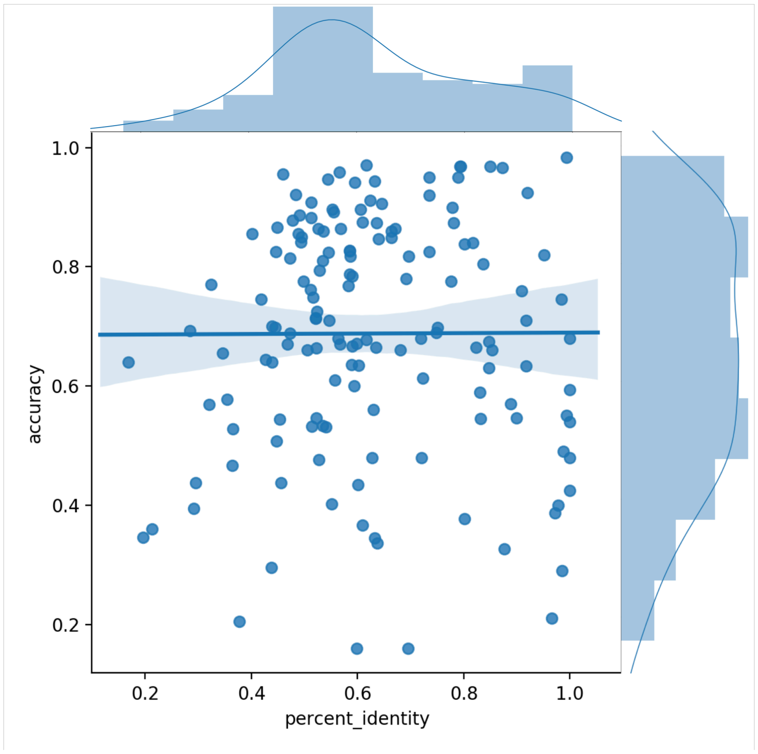


**SI Fig. 5** – The average DNA percent identity (x axis) and oxidoreductase classifier model accuracy (y axis) for genes within each of the EC annotations in the *oxidoreductase model* validation set. Each dot represents a unique EC number.

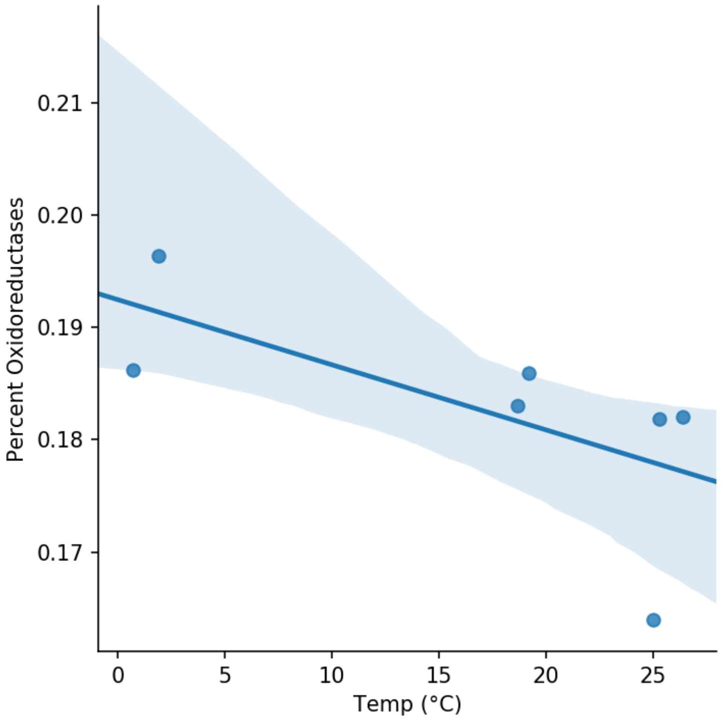


**SI Fig. 6** – The proportion of reads identified as oxidoreductases by the oxidoreductase classifier (y axis), correlated with temperature (x axis), in surface water metagenomes from the *oxidoreductase metagenome set*. $R^2$ = -0.66, P=0.11.

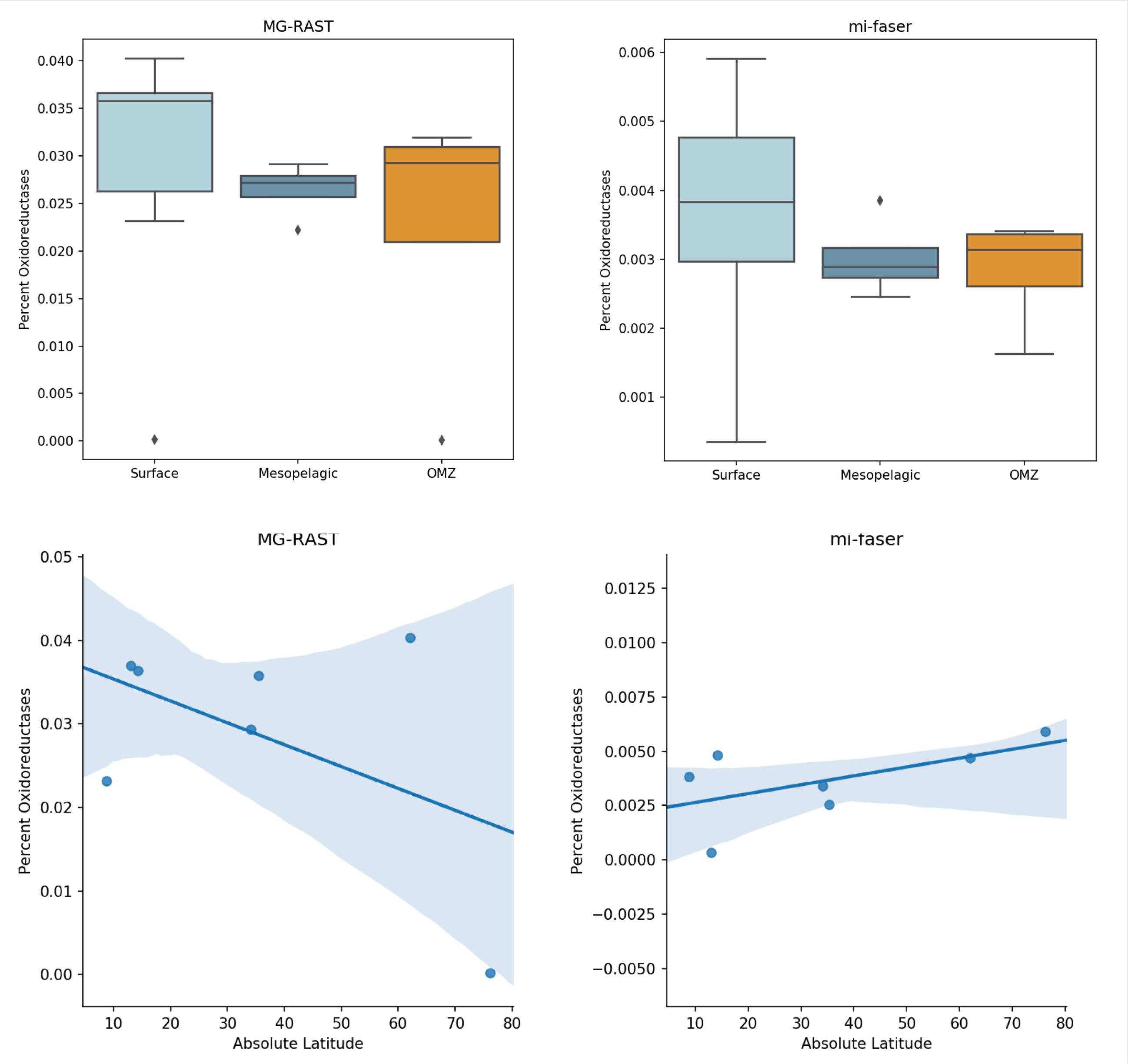


**SI Fig. 7** – Trends in the proportion of oxidoreductases predicted in metagenomes from the *oxidoreductase metagenome set* for functional annotations from MG-RAST (left column) and mi-faser (right column). There are no significant differences in the proportion of oxidoreductases with depth (top row, ANOVA P=0.73 for MG-RAST, P=0.60 for mi-faser), and no significant trends in surface waters with latitude (bottom row, $R^2$= -0.49 P=0.27 for MG-RAST, and $R^2$=0.58 P=0.17 for mi-faser).

### Supplemental Tables

| GenBank Assembly Accession | Assembly Name | BioSample ID | NCBI Organism Name | Genome Accession | dataset |
|---|---|---|---|---|---|
| GCA_000526435.1 | ASM52643v1 | SAMN02584936 | Caldicoprobacter oshimai DSM 21659 | GCF_000526435.1 | train |
| GCA_000166775.1 | ASM16677v1 | SAMN00713565 | Caldicellulosiruptor kronotskyensis 2002 | GCF_000166775.1 | train |
| GCA_000328765.2 | ASM32876v2 | SAMEA2272437 | Tepidanaerobacter acetatoxydans Re1 | GCF_000328765.2 | train |
| GCA_000499205.1 | MAEPY2 1.0 | SAMN02343193 | Paenibacillus sp. MAEPY2 | GCF_000499205.1 | train |
| GCA_001730225.1 | ASM173022v1 | SAMN05731213 | Desulfuribacillus alkaliarsenatis | GCF_001730225.1 | train |
| GCA_001552655.1 | ASM155265v1 | SAMD00045739 | Alicyclobacillus kakegawensis NBRC 103104 | GCF_001552655.1 | train |
| none | none | none | none | GCA_003456095.1 | train |
| GCA_000020005.1 | ASM2000v1 | SAMN02598430 | Natranaerobius thermophilus JW/NM-WN-LF | GCF_000020005.1 | train |
| GCA_900111575.1 | IMG-taxon 2602042032 annotated assembly | SAMN03080614 | Anaerobranca gottschalkii DSM 13577 | GCF_900111575.1 | train |
| GCA_002427055.1 | ASM242705v1 | SAMN06457451 | Firmicutes bacterium UBA5500 | GCA_002427055.1 | train |
| GCA_002919235.1 | ASM291923v1 | SAMN08158218 | Clostridia bacterium | GCA_002919235.1 | train |
| GCA_002399235.1 | ASM239923v1 | SAMN06451266 | Firmicutes bacterium UBA4881 | GCA_002399235.1 | train |
| GCA_002408345.1 | ASM240834v1 | SAMN06453845 | Firmicutes bacterium UBA5301 | GCA_002408345.1 | train |
| GCA_002426275.1 | ASM242627v1 | SAMN06454099 | Firmicutes bacterium UBA5499 | GCA_002426275.1 | train |
| GCA_002452295.1 | ASM245229v1 | SAMN06456985 | Firmicutes bacterium UBA6811 | GCA_002452295.1 | train |
| GCA_900016865.1 | Clostridia bin genome 5 | SAMEA3730008 | uncultured Clostridia bacterium | GCA_900016865.1 | train |
| GCA_002426645.1 | ASM242664v1 | SAMN06456060 | Firmicutes bacterium UBA5435 | GCA_002426645.1 | train |
| none | none | none | none | GCA_003501335.1 | train |
| GCA_003023725.1 | ASM302372v1 | SAMN08683243 | Sulfobacillus benefaciens | GCA_003023725.1 | train |
| GCA_000009905.1 | ASM990v1 | SAMD00061067 | Symbiobacterium thermophilum IAM 14863 | GCF_000009905.1 | train |
| GCA_002919105.1 | ASM291910v1 | SAMN08158229 | Thermaerobacter sp. | GCA_002919105.1 | train |
| none | none | none | none | GCA_003517845.1 | train |
| GCA_002375925.1 | ASM237592v1 | SAMN06455820 | Firmicutes bacterium UBA3575 | GCA_002375925.1 | train |
| GCA_001513125.1 | ASM151312v1 | SAMN03778954 | Clostridia bacterium DTU030 | GCA_001513125.1 | train |
| GCA_900104055.1 | IMG-taxon 2623620517 annotated assembly | SAMN05216366 | Selenomonas ruminantium | GCF_900104055.1 | train |
| GCA_000224515.2 | ASM22451v1 | SAMN02471585 | Desulfosporosinus sp. OT | GCF_000224515.1 | train |
| GCA_001512665.1 | ASM151266v1 | SAMN03776771 | Clostridiales bacterium DTU073 | GCA_001512665.1 | train |
| GCA_001875325.1 | ASM187532v1 | SAMN05430241 | Moorella thermoacetica | GCF_001875325.1 | train |
| GCA_002418965.1 | ASM241896v1 | SAMN06454537 | Peptococcaceae bacterium UBA5757 | GCA_002418965.1 | train |
| GCA_002418765.1 | ASM241876v1 | SAMN06451449 | Peptococcaceae bacterium UBA5767 | GCA_002418765.1 | train |
| GCA_002840245.1 | ASM284024v1 | SAMN06767695 | Firmicutes bacterium HGW-Firmicutes-15 | GCA_002840245.1 | train |
| GCA_000016165.1 | ASM1616v1 | SAMN02598304 | Desulfotomaculum reducens MI-1 | GCF_000016165.1 | train |
| GCA_002840165.1 | ASM284016v1 | SAMN06767708 | Firmicutes bacterium HGW-Firmicutes-8 | GCA_002840165.1 | train |
| GCA_003054495.1 | ASM305449v1 | SAMN07757920 | Carboxydocella thermautotrophica | GCF_003054495.1 | train |
| GCA_900111505.1 | IMG-taxon 2642422559 annotated assembly | SAMN04515653 | Halanaerobium congolense | GCF_900111505.1 | train |
| GCA_000187935.2 | ASM18793v2 | SAMN02436555 | Streptococcus parauberis NCFD 2020 | GCF_000187935.1 | train |
| GCA_000622245.1 | ASM62224v1 | SAMN02743999 | Fusobacterium perfoetens ATCC 29250 | GCF_000622245.1 | train |
| GCA_000012505.1 | ASM1250v1 | SAMN02598312 | Synechococcus sp. CC9902 | GCF_000012505.1 | train |
| GCA_900313115.1 | Rumen uncultured genome RUG770 | SAMEA104666321 | uncultured bacterium | GCA_900313115.1 | train |
| GCA_002083825.1 | ASM208382v1 | SAMN05981169 | Candidatus Sericytochromatia bacterium S15B-MN24 CBMW_12 | GCA_002083825.1 | train |
| GCA_001771545.1 | ASM177154v1 | SAMN04314412 | candidate division WOR-1 bacterium RIFOXYB2_FULL_42_35 | GCA_001771545.1 | train |
| GCA_003265885.1 | ASM326588v1 | SAMN08965200 | Candidatus Marinamargulisbacteria bacterium SCGC AG-343-D04 | GCA_003265885.1 | train |
| GCA_003242895.1 | ASM324289v1 | SAMN09222456 | Candidatus Margulisbacteria bacterium | GCA_003242895.1 | train |
| GCA_002413305.1 | ASM241330v1 | SAMN06451868 | Actinobacteria bacterium UBA5176 | GCA_002413305.1 | train |
| GCA_003138855.1 | 20110800_S2D | SAMN08179187 | Acidimicrobiaceae bacterium | GCA_003138855.1 | train |
| GCA_002897715.1 | ASM289771v1 | SAMD00081361 | bacterium BMS3Abin01 | GCA_002897715.1 | train |
| GCA_000014185.1 | ASM1418v1 | SAMN02598258 | Rubrobacter xylanophilus DSM 9941 | GCF_000014185.1 | train |
| GCA_002331575.1 | ASM233157v1 | SAMN06454938 | Atopobium sp. UBA2090 | GCA_002331575.1 | train |
| GCA_002366545.1 | ASM236654v1 | SAMN06453901 | Actinobacteria bacterium UBA3085 | GCA_002366545.1 | train |
| GCA_002779205.1 | ASM277920v1 | SAMN06659258 | Actinobacteria bacterium CG08_land_8_20_14_0_20_35_9 | GCA_002779205.1 | train |
| GCA_003157385.1 | 20120600_E2D | SAMN08179582 | Actinobacteria bacterium | GCA_003157385.1 | train |
| GCA_003133745.1 | 20120500_P13 | SAMN08179405 | candidate division WPS-2 bacterium | GCA_003133745.1 | train |
| GCA_002423485.1 | ASM242348v1 | SAMN06454064 | bacterium UBP9_UBA6111 | GCA_002423485.1 | train |
| GCA_001443485.1 | ASM144348v1 | SAMN03462097 | Armatimonadetes bacterium CSP1-3 | GCA_001443485.1 | train |
| GCA_002254605.1 | ASM225460v1 | SAMN07230129 | Armatimonadetes bacterium JP3_11 | GCA_002254605.1 | train |
| GCA_002898895.1 | ASM289889v1 | SAMD00093766 | bacterium HR16 | GCA_002898895.1 | train |
| GCA_000427095.1 | T49 | SAMEA2272150 | Chthonomonas calidirosea T49 | GCF_000427095.1 | train |
| GCA_002431715.1 | ASM243171v1 | SAMN06453966 | Armatimonadetes bacterium UBA5829 | GCA_002431715.1 | train |
| GCA_003134215.1 | 20111000_S2S | SAMN08179378 | Armatimonadetes bacterium | GCA_003134215.1 | train |
| GCA_001872605.1 | ASM187260v1 | SAMN04328248 | Armatimonadetes bacterium CG2_30_59_28 | GCA_001872605.1 | train |
| GCA_002898575.1 | ASM289857v1 | SAMD00093767 | bacterium HR17 | GCA_002898575.1 | train |
| GCA_002409445.1 | ASM240944v1 | SAMN06456318 | Armatimonadetes bacterium UBA5419 | GCA_002409445.1 | train |
| GCA_002411015.1 | ASM241101v1 | SAMN06451880 | Armatimonadetes bacterium UBA5352 | GCA_002411015.1 | train |
| GCA_002410925.1 | ASM241092v1 | SAMN06451889 | bacterium UBP13_UBA5359 | GCA_002410925.1 | train |
| GCA_001003605.1 | ASM100360v1 | SAMN03319960 | Parcubacteria group bacterium GW2011_GWC2_48_17 | GCA_001003605.1 | train |
| GCA_002328565.1 | ASM232856v1 | SAMN06453241 | Candidatus Moranbacteria bacterium UBA2206 | GCA_002328565.1 | train |
| GCA_002792135.1 | ASM279213v1 | SAMN06659773 | bacterium (Candidatus Torokbacteria) CG_4_10_14_0_2_um_filter_35_8 | GCA_002792135.1 | train |
| GCA_001791275.1 | ASM179127v1 | SAMN04315591 | Candidatus Uhrbacteria bacterium RIFCSPHIGHO2_12_FULL_60_25 | GCA_001791275.1 | train |
| GCA_002773695.1 | ASM277369v1 | SAMN06659340 | Candidatus Doudnabacteria bacterium CG10_big_fil_rev_8_21_14_0_10_41_10 | GCA_002773695.1 | train |
| GCA_002686445.1 | ASM268644v1 | SAMN07618373 | bacterium | GCA_002686445.1 | train |
| GCA_000999315.1 | ASM99931v1 | SAMN03319709 | Candidatus Peregrinibacteria bacterium GW2011_GWF2_43_17 | GCA_000999315.1 | train |
| GCA_002482925.1 | ASM248292v1 | SAMN06452856 | Candidatus Saccharibacteria bacterium UBA7683 | GCA_002482925.1 | train |
| GCA_001872945.1 | ASM187294v1 | SAMN04328267 | bacterium CG2_30_37_16 | GCA_001872945.1 | train |
| GCA_002792735.1 | ASM279273v1 | SAMN06659745 | bacterium CG_4_10_14_0_2_um_filter_33_32 | GCA_002792735.1 | train |
| GCA_002780235.1 | ASM278023v1 | SAMN06659215 | Candidatus Berkelbacteria bacterium CG06_land_8_20_14_3_00_43_10 | GCA_002780235.1 | train |
| GCA_001771185.1 | ASM177118v1 | SAMN04316212 | candidate division Kazan bacterium RIFCSPHIGHO2_01_FULL_44_14 | GCA_001771185.1 | train |
| GCA_001773695.1 | ASM177369v1 | SAMN04315142 | Candidatus Amesbacteria bacterium RIFCSPLOWO2_01_FULL_48_25 | GCA_001773695.1 | train |
| GCA_001772615.1 | ASM177261v1 | SAMN04314869 | candidate division WWE3 bacterium RIFCSPHIGHO2_01_FULL_40_23 | GCA_001772615.1 | train |
| GCA_001771135.1 | ASM177113v1 | SAMN04313865 | candidate division CPR3 bacterium GWF2_35_18 | GCA_001771135.1 | train |
| GCA_002771355.1 | ASM277135v1 | SAMN06659655 | candidate division WWE3 bacterium CG22_combo_CG10-13_8_21_14_all_39_12 | GCA_002771355.1 | train |
| GCA_001567305.1 | ASM156730v1 | SAMN03372540 | candidate division WS6 bacterium OLB21 | GCA_001567305.1 | train |
| GCA_001873755.1 | ASM187375v1 | SAMN04328296 | Candidatus Wirthbacteria bacterium CG2_30_54_11 | GCA_001873755.1 | train |
| GCA_002311865.1 | ASM231186v1 | SAMN06457163 | Dehalococcoidia bacterium UBA1151 | GCA_002311865.1 | train |
| GCA_002436065.1 | ASM243606v1 | SAMN06455613 | Chloroflexi bacterium UBA6077 | GCA_002436065.1 | train |
| GCA_003158355.1 | 20120600_E3D | SAMN08179596 | Anaerolineaceae bacterium | GCA_003158355.1 | train |
| GCA_002532535.1 | ASM253253v1 | SAMN07514207 | Chloroflexi bacterium Kir15-3F | GCF_002532535.1 | train |
| GCA_003268475.1 | ASM326847v1 | SAMN04600100 | Thermogemmatispora tikiterensis | GCA_003268475.1 | train |
| GCA_002413265.1 | ASM241326v1 | SAMN06451869 | Chloroflexi bacterium UBA5177 | GCA_002413265.1 | train |
| GCA_002347925.1 | ASM234792v1 | SAMN06453796 | Chloroflexi bacterium UBA2235 | GCA_002347925.1 | train |
| GCA_002404055.1 | ASM240405v1 | SAMN06451262 | Chloroflexi bacterium UBA4733 | GCA_002404055.1 | train |
| GCA_003135255.1 | 20111000_S2D | SAMN08179358 | candidate division AD3 bacterium | GCA_003135255.1 | train |
| GCA_003157875.1 | 20120600_E1D | SAMN08179544 | Chloroflexi bacterium | GCA_003157875.1 | train |
| GCA_002433065.1 | ASM243306v1 | SAMN06452350 | Chloroflexi bacterium UBA6622 | GCA_002433065.1 | train |
| GCA_002355995.1 | ASM235599v1 | SAMD00066626 | Thermus thermophilus | GCF_002355995.1 | train |
| GCA_002050035.1 | ASM205003v1 | SAMN06287783 | Planctomycetales bacterium 4484_113 | GCA_002050035.1 | train |
| GCA_002435745.1 | ASM243574v1 | SAMN06453690 | bacterium UBP15_UBA6099 | GCA_002435745.1 | train |
| GCA_001644665.1 | ASM164466v1 | SAMN02954477 | Fervidobacterium pennivorans | GCF_001644665.1 | train |
| GCA_900465355.1 | Ran1-1 | SAMEA4698766 | Candidatus Bipolaricaulis anaerobius | GCA_900465355.1 | train |
| GCA_002877955.1 | ASM287795v1 | SAMN08107316 | Caldisericum exile | GCA_002877955.1 | train |
| none | none | none | none | GCA_003476705.1 | train |
| GCA_002878375.1 | ASM287837v1 | SAMN07140561 | Dictyoglomus sp. NZ13-RE01 | GCA_002878375.1 | train |
| GCA_003057965.1 | ASM305796v1 | SAMN06013554 | Thermodesulfobium acidiphilum | GCF_003057965.1 | train |
| GCA_001604435.1 | ASM160443v1 | SAMN03837571 | Synergistales bacterium Syner_01 | GCA_001604435.1 | train |
| GCA_001508835.1 | ASM150883v1 | SAMN03445127 | bacterium 42_11 | GCA_001508835.1 | train |
| GCA_000405345.1 | ASM40534v1 | SAMN02441602 | Atribacteria bacterium SCGC AAA255-N14 | GCA_000405345.1 | train |
| GCA_000353875.1 | "Ca. Caldatribacterium californiense" | SAMN01797434 | Candidatus Caldatribacterium californiense OP9-cSCG | GCA_000353875.1 | train |
| GCA_001872985.1 | ASM187298v1 | SAMN04328278 | bacterium CG2_30_54_10 | GCA_001872985.1 | train |
| GCA_002422125.1 | ASM242212v1 | SAMN06457399 | bacterium UBP17_UBA6191 | GCA_002422125.1 | train |
| GCA_002869225.1 | ASM286922v1 | SAMN07982665 | bacterium BM706 | GCA_002869225.1 | train |
| GCA_001791795.1 | ASM179179v1 | SAMN04314129 | Candidatus Wallbacteria bacterium GWC2_49_35 | GCA_001791795.1 | train |
| GCA_900105685.1 | IMG-taxon 2690315640 annotated assembly | SAMN05444156 | Verrucomicrobium sp. GAS474 | GCF_900105685.1 | train |
| GCA_900320225.1 | Rumen uncultured genome RUG572 | SAMEA104666872 | uncultured Lentisphaerae bacterium | GCA_900320225.1 | train |
| GCA_001803205.1 | ASM180320v1 | SAMN04313868 | Lentisphaerae bacterium GWF2_38_69 | GCA_001803205.1 | train |
| GCA_002308555.1 | ASM230855v1 | SAMN06455243 | bacterium UBP3_UBA1247 | GCA_002308555.1 | train |
| GCA_002329605.1 | ASM232960v1 | SAMN06451854 | bacterium UBP3_UBA1439 | GCA_002329605.1 | train |
| GCA_002895085.1 | ASM289508v1 | SAMN07767759 | Chlamydia abortus | GCF_002895085.1 | train |
| GCA_000338295.1 | First | SAMN02261079 | Rhodopirellula europaea 6C | GCF_000338295.1 | train |
| GCA_002364005.1 | ASM236400v1 | SAMN06451765 | Phycisphaerales bacterium UBA3190 | GCA_002364005.1 | train |
| GCA_003170085.1 | 20110800_E3D | SAMN08179086 | Planctomycetia bacterium | GCA_003170085.1 | train |
| GCA_001302825.1 | ASM130282v1 | SAMN03994142 | Planctomycetes bacterium DG_23 | GCA_001302825.1 | train |
| GCA_002311925.1 | ASM231192v1 | SAMN06457040 | Planctomycetes bacterium UBA1135 | GCA_002311925.1 | train |
| GCA_002687715.1 | ASM268771v1 | SAMN07619727 | Planctomycetes bacterium | GCA_002687715.1 | train |
| GCA_002320775.1 | ASM232077v1 | SAMN06451693 | Planctomycetes bacterium UBA1662 | GCA_002320775.1 | train |
| GCA_002343805.1 | ASM234380v1 | SAMN06454447 | Planctomycetes bacterium UBA2392 | GCA_002343805.1 | train |
| none | none | none | none | GCA_003482625.1 | train |
| GCA_003249795.1 | ASM324979v1 | SAMN09288115 | bacterium | GCA_003249795.1 | train |
| GCA_002393835.1 | ASM239383v1 | SAMN06453632 | Treponema sp. UBA4326 | GCA_002393835.1 | train |
| GCA_001829125.1 | ASM182912v1 | SAMN04314119 | Spirochaetes bacterium GWB1_27_13 | GCA_001829125.1 | train |
| GCA_001829155.1 | ASM182915v1 | SAMN04313962 | Spirochaetes bacterium GWB1_36_13 | GCA_001829155.1 | train |
| GCA_900112165.1 | IMG-taxon 2597490362 annotated assembly | SAMN02745150 | Brevinema andersonii | GCF_900112165.1 | train |
| GCA_001746225.1 | ASM174622v1 | SAMN05209828 | Brachyspira hampsonii | GCF_001746225.1 | train |
| none | none | none | none | GCA_003497555.1 | train |
| GCA_002786225.1 | ASM278622v1 | SAMN07572185 | Leptospira sp. | GCA_002786225.1 | train |
| GCA_002839285.1 | ASM283928v1 | SAMN06767754 | Spirochaetae bacterium HGW-Spirochaetae-5 | GCA_002839285.1 | train |
| GCA_002716355.1 | ASM271635v1 | SAMN07618404 | bacterium | GCA_002716355.1 | train |
| GCA_001650905.1 | ASM165090v1 | SAMN04957308 | Balneola sp. EhC07 | GCF_001650905.1 | train |
| GCA_000020465.1 | ASM2046v1 | SAMN02598270 | Chlorobium limicola DSM 245 | GCF_000020465.1 | train |
| none | none | none | none | GCA_003538625.1 | train |
| GCA_003142475.1 | 20120800_E3D | SAMN08180131 | Ignavibacteriae bacterium | GCA_003142475.1 | train |
| GCA_001536065.1 | JGI assembly | SAMN03436147 | Candidatus Chrysopegis kryptomonas | GCF_001536065.1 | train |
| GCA_002418685.1 | ASM241868v1 | SAMN06455695 | Ignavibacteria bacterium UBA5771 | GCA_002418685.1 | train |
| GCA_003246455.1 | ASM324645v1 | SAMN09287913 | Bacteroidetes bacterium | GCA_003246455.1 | train |
| GCA_002327255.1 | ASM232725v1 | SAMN06452215 | bacterium UBA2214 | GCA_002327255.1 | train |
| GCA_001304035.1 | ASM130403v1 | SAMN03994197 | bacterium SM23_31 | GCA_001304035.1 | train |
| GCA_001886815.1 | ASM188681v1 | SAMN05946030 | Caldithrix abyssi DSM 13497 | GCF_001886815.1 | train |
| GCA_002726775.1 | ASM272677v1 | SAMN07619376 | Candidatus Marinimicrobia bacterium | GCA_002726775.1 | train |
| GCA_002452555.1 | ASM245255v1 | SAMN06456991 | Candidatus Marinimicrobia bacterium UBA6817 | GCA_002452555.1 | train |
| GCA_001872685.1 | ASM187268v1 | SAMN04328203 | Candidatus Marinimicrobia bacterium CG1_02_48_14 | GCA_001872685.1 | train |
| GCA_002774355.1 | ASM277435v1 | SAMN06659292 | Candidatus Marinimicrobia bacterium CG08_land_8_20_14_0_20_45_22 | GCA_002774355.1 | train |
| GCA_002382735.1 | ASM238273v1 | SAMN06454802 | bacterium UBP11_UBA4055 | GCA_002382735.1 | train |
| GCA_002898195.1 | ASM289819v1 | SAMD00081381 | bacterium BMS3Bbin04 | GCA_002898195.1 | train |
| GCA_900177705.1 | IMG-taxon 2718218506 annotated assembly | SAMN05720473 | Fibrobacter sp. UWB15 | GCF_900177705.1 | train |
| GCA_001789205.1 | ASM178920v1 | SAMN04314369 | Candidatus Raymondbacteria bacterium RIFOXYA2_FULL_49_16 | GCA_001789205.1 | train |
| GCA_001462255.1 | ASM146225v1 | SAMN03999395 | Fibrobacteria bacterium GUT77 | GCA_001462255.1 | train |
| GCA_002435865.1 | ASM243586v1 | SAMN06453688 | Cloacimonetes bacterium UBA6097 | GCA_002435865.1 | train |
| GCA_003222155.1 | ASM322215v1 | SAMN08911877 | Gemmatimonadetes bacterium | GCA_003222155.1 | train |
| GCA_001780825.1 | ASM178082v1 | SAMN04315771 | Candidatus Glassbacteria bacterium RIFCSPLOWO2_12_FULL_58_11 | GCA_001780825.1 | train |
| GCA_002403075.1 | ASM240307v1 | SAMN06455476 | bacterium UBP1_UBA4783 | GCA_002403075.1 | train |
| GCA_001780165.1 | ASM178016v1 | SAMN04313721 | Candidatus Eisenbacteria bacterium RBG_16_71_46 | GCA_001780165.1 | train |
| GCA_002686915.1 | ASM268691v1 | SAMN07619151 | Gemmatimonadetes bacterium | GCA_002686915.1 | train |
| GCA_002686955.1 | ASM268695v1 | SAMN07619150 | Gemmatimonadetes bacterium | GCA_002686955.1 | train |
| GCA_001777915.1 | ASM177791v1 | SAMN04314394 | Candidatus Edwardsbacteria bacterium RifOxyA12_full_54_48 | GCA_001777915.1 | train |
| GCA_001303705.1 | ASM130370v1 | SAMN03994202 | candidate division TA06 bacterium SM23_40 | GCA_001303705.1 | train |
| GCA_001304015.1 | ASM130401v1 | SAMN03994216 | candidate division Zixibacteria bacterium SM23_81 | GCA_001304015.1 | train |
| GCA_002084765.1 | ASM208476v1 | SAMN06638386 | Candidatus Cloacimonetes bacterium 4572_55 | GCA_002084765.1 | train |
| GCA_002712305.1 | ASM271230v1 | SAMN07619168 | Gemmatimonadetes bacterium | GCA_002712305.1 | train |
| GCA_002049985.1 | ASM204998v1 | SAMN06288098 | Candidatus Latescibacteria bacterium 4484_181 | GCA_002049985.1 | train |
| GCA_002059205.1 | ASM205920v1 | SAMN06287578 | Candidatus Latescibacteria bacterium 4484_107 | GCA_002059205.1 | train |
| GCA_000403035.1 | ASM40303v1 | SAMN02441742 | Candidatus Latescibacter anaerobius SCGC AAA252-E07 | GCA_000403035.1 | train |
| GCA_002085375.1 | ASM208537v1 | SAMN06624697 | candidate division Zixibacteria bacterium 4484_95 | GCA_002085375.1 | train |
| GCA_002085385.1 | ASM208538v1 | SAMN06624686 | candidate division Zixibacteria bacterium 4484_93 | GCA_002085385.1 | train |
| GCA_002316275.1 | ASM231627v1 | SAMN06455730 | candidate division WOR-3 bacterium UBA1063 | GCA_002316275.1 | train |
| GCA_000494225.1 | Calescamantes bacterium OTU 1 | SAMN02441764 | Candidatus Calescibacterium nevadense OTU 1 | GCA_000494225.1 | train |
| GCA_001303785.1 | ASM130378v1 | SAMN03994203 | candidate division WOR_3 bacterium SM23_42 | GCA_001303785.1 | train |
| GCA_002366725.1 | ASM236672v1 | SAMN06455134 | candidate division WOR-3 bacterium UBA3073 | GCA_002366725.1 | train |
| GCA_002256535.1 | ASM225653v1 | SAMN07350899 | candidate division bacterium WOR-3 4484_18 | GCA_002256535.1 | train |
| GCA_002215665.1 | ASM221566v1 | SAMN06887667 | candidate division TA06 bacterium JGI_Cruoil_03_38_101 | GCA_002215665.1 | train |
| GCA_001508335.1 | ASM150833v1 | SAMN03445151 | candidate division TA06 bacterium 32_111 | GCA_001508335.1 | train |
| GCA_001412365.1 | ASM141236v1 | SAMN04125646 | Desulfuromonas sp. SDB | GCA_001412365.1 | train |
| GCA_002030045.1 | ASM203004v1 | SAMN04330451 | Candidatus Aegiribacteria bacterium MLS_C | GCA_002030045.1 | train |
| GCA_002699335.1 | ASM269933v1 | SAMN07619758 | Candidatus Poribacteria bacterium | GCA_002699335.1 | train |
| GCA_002746185.1 | ASM274618v1 | SAMN07568857 | Candidatus Hydrogenedentes bacterium | GCA_002746185.1 | train |
| GCA_001567115.1 | ASM156711v1 | SAMN03372536 | Omnitrophica bacterium OLB16 | GCA_001567115.1 | train |
| GCA_001774505.1 | ASM177450v1 | SAMN04313985 | Candidatus Coatesbacteria bacterium RBG_13_66_14 | GCA_001774505.1 | train |
| GCA_002842095.1 | ASM284209v1 | SAMN06767623 | candidate division BRC1 bacterium HGW-BRC1-1 | GCA_002842095.1 | train |
| GCA_002715745.1 | ASM271574v1 | SAMN07620247 | Woeseiaceae bacterium | GCA_002715745.1 | train |
| GCA_002779605.1 | ASM277960v1 | SAMN06659231 | Zetaproteobacteria bacterium CG06_land_8_20_14_3_00_59_53 | GCA_002779605.1 | train |
| GCA_002109495.1 | ASM210949v1 | SAMN04574817 | Magnetofaba australis IT-1 | GCF_002109495.1 | train |
| GCA_000992305.1 | ASM99230v1 | SAMN03319398 | candidate division TM6 bacterium GW2011_GWF2_38_10 | GCA_000992305.1 | train |
| GCA_002325765.1 | ASM232576v1 | SAMN06456904 | bacterium UBA1459 | GCA_002325765.1 | train |
| GCA_002309575.1 | ASM230957v1 | SAMN06454758 | bacterium UBP6_UBA1209 | GCA_002309575.1 | train |
| GCA_002082305.1 | ASM208230v1 | SAMN04448864 | SAR324 cluster bacterium SAR324-CTD7B | GCA_002082305.1 | train |
| GCA_002913305.1 | ASM291330v1 | SAMN08200415 | Campylobacter concisus | GCF_002913305.1 | train |
| GCA_002119425.1 | ASM211942v1 | SAMN05511439 | Desulfurella amilsii | GCF_002119425.1 | train |
| GCA_002878035.1 | ASM287803v1 | SAMN08107306 | Sulfurihydrogenibium sp. | GCA_002878035.1 | train |
| GCA_000421485.1 | ASM42148v1 | SAMN02440891 | Desulfurobacterium sp. TC5-1 | GCF_000421485.1 | train |
| GCA_001547735.1 | ASM154773v1 | SAMD00061062 | Thermosulfidibacter takaii ABI70S6 | GCF_001547735.1 | train |
| GCA_002878065.1 | ASM287806v1 | SAMN08107314 | Calditerrivibrio nitroreducens | GCA_002878065.1 | train |
| GCA_000469585.1 | Chrysiogenes arsenatis | SAMN02471296 | Chrysiogenes arsenatis DSM 11915 | GCF_000469585.1 | train |
| GCA_002708105.1 | ASM270810v1 | SAMN07618396 | bacterium | GCA_002708105.1 | train |
| GCA_003232385.1 | ASM323238v1 | SAMN09240063 | bacterium | GCA_003232385.1 | train |
| GCA_003252055.1 | ASM325205v1 | SAMN09288001 | Desulfobacter postgatei | GCA_003252055.1 | train |
| GCA_001603845.1 | ASM160384v1 | SAMN03837530 | Syntrophobacterales bacterium Delta_01 | GCA_001603845.1 | train |
| GCA_002011835.1 | ASM201183v1 | SAMN06226412 | Desulfarculaceae bacterium JdFR-95 | GCA_002011835.1 | train |
| GCA_000195295.1 | ASM19529v1 | SAMN00713603 | Desulfobacca acetoxidans DSM 11109 | GCF_000195295.1 | train |
| GCA_001797675.1 | ASM179767v1 | SAMN04314000 | Deltaproteobacteria bacterium RBG_13_43_22 | GCA_001797675.1 | train |
| none | none | none | none | GCA_003509305.1 | train |
| GCA_000423845.1 | ASM42384v1 | SAMN02441489 | Thermodesulfobacterium hveragerdense DSM 12571 | GCF_000423845.1 | train |
| GCA_001687335.1 | ASM168733v1 | SAMN05251087 | Dissulfuribacter thermophilus | GCF_001687335.1 | train |
| GCA_001304365.1 | ASM130436v1 | SAMN03994151 | Syntrophobacter sp. DG_60 | GCA_001304365.1 | train |
| GCA_003142535.1 | 20120800_E2X | SAMN08180125 | Syntrophaceae bacterium | GCA_003142535.1 | train |
| GCA_002010915.1 | ASM201091v1 | SAMN06226414 | Desulfarculaceae bacterium JdFR-97 | GCA_002010915.1 | train |
| GCA_001797905.1 | ASM179790v1 | SAMN04314029 | Deltaproteobacteria bacterium RBG_16_54_11 | GCA_001797905.1 | train |
| GCA_002418945.1 | ASM241894v1 | SAMN06454534 | Deltaproteobacteria bacterium UBA5758 | GCA_002418945.1 | train |
| GCA_002316295.1 | ASM231629v1 | SAMN06455729 | Syntrophaceae bacterium UBA1062 | GCA_002316295.1 | train |
| GCA_002841765.1 | ASM284176v1 | SAMN06767655 | Deltaproteobacteria bacterium HGW-Deltaproteobacteria-4 | GCA_002841765.1 | train |
| GCA_001795535.1 | ASM179553v1 | SAMN04314173 | Deltaproteobacteria bacterium GWA2_54_12 | GCA_001795535.1 | train |
| GCA_002839535.1 | ASM283953v1 | SAMN06767742 | Nitrospira bacterium HGW-Nitrospira-1 | GCA_002839535.1 | train |
| GCA_001458695.1 | NiCh1 | SAMEA3560316 | Candidatus Nitrospira inopinata | GCF_001458695.1 | train |
| GCA_001803705.1 | ASM180370v1 | SAMN04313879 | Nitrospirae bacterium GWC2_56_14 | GCA_001803705.1 | train |
| GCA_001803795.1 | ASM180379v1 | SAMN04313704 | Nitrospirae bacterium RBG_16_64_22 | GCA_001803795.1 | train |
| GCA_001873285.1 | ASM187328v1 | SAMN04328285 | Nitrospirae bacterium CG2_30_53_67 | GCA_001873285.1 | train |
| GCA_003221515.1 | ASM322151v1 | SAMN08911940 | Candidatus Rokubacteria bacterium | GCA_003221515.1 | train |
| GCA_000739515.1 | ASM73951v1 | SAMD00016904 | Candidatus Moduliflexus flocculans | GCA_000739515.1 | train |
| GCA_003230635.1 | ASM323063v1 | SAMN09240166 | bacterium | GCA_003230635.1 | train |
| GCA_002731985.1 | ASM273198v1 | SAMN07619521 | Nitrospinae bacterium | GCA_002731985.1 | train |
| GCA_002500605.1 | ASM250060v1 | SAMN06450621 | Nitrospinaceae bacterium UBA7883 | GCA_002500605.1 | train |
| GCA_001804915.1 | ASM180491v1 | SAMN04315473 | Nitrospinae bacterium RIFCSPHIGHO2_02_FULL_39_82 | GCA_001804915.1 | train |
| GCA_001803565.1 | ASM180356v1 | SAMN04315881 | Nitrospinae bacterium RIFCSPLOWO2_12_FULL_45_22 | GCA_001803565.1 | train |
| GCA_002377645.1 | ASM237764v1 | SAMN06457116 | Nitrospinaceae bacterium UBA3496 | GCA_002377645.1 | train |
| GCA_000522425.1 | v3 | SAMN02420412 | Candidatus Entotheonella factor | GCA_000522425.1 | train |
| GCA_001790715.1 | ASM179071v1 | SAMN04314270 | Candidatus Schekmanbacteria bacterium RBG_16_38_11 | GCA_001790715.1 | train |
| GCA_003228585.1 | ASM322858v1 | SAMN09240188 | Proteobacteria bacterium | GCA_003228585.1 | train |
| GCA_001798995.1 | ASM179899v1 | SAMN04315407 | Deltaproteobacteria bacterium RIFCSPHIGHO2_02_FULL_60_17 | GCA_001798995.1 | train |
| GCA_000526155.1 | ASM52615v1 | SAMN02584914 | Deferrisoma camini S3R1 | GCA_000526155.1 | train |
| GCA_001797445.1 | ASM179744v1 | SAMN04313911 | Deltaproteobacteria bacterium GWA2_65_63 | GCA_001797445.1 | train |
| GCA_003170555.1 | 20110800_E3D | SAMN08179068 | Deltaproteobacteria bacterium | GCA_003170555.1 | train |

| | | | | | |
|---|---|---|---|---|---|
| GCA_001873295.1 | ASM187329v1 | SAMN04328286 | Nitrospirae bacterium CG2_30_70_394 | GCA_001873295.1 | train |
| GCA_001797815.1 | ASM179781v1 | SAMN04313688 | Deltaproteobacteria bacterium RBG_13_61_14 | GCA_001797815.1 | train |
| GCA_003153015.1 | 20120700_E3M | SAMN08179787 | Myxococcales bacterium | GCA_003153015.1 | train |
| GCA_002309695.1 | ASM230969v1 | SAMN06454754 | Myxococcales bacterium UBA1203 | GCA_002309695.1 | train |
| GCA_001798605.1 | ASM179860v1 | SAMN04314410 | Deltaproteobacteria bacterium RIFOXYA12_FULL_58_15 | GCA_001798605.1 | train |
| GCA_002722845.1 | ASM272284v1 | SAMN07619499 | Myxococcales bacterium | GCA_002722845.1 | train |
| GCA_003258315.1 | ASM325831v1 | SAMN09396997 | Bradymonas sediminis | GCF_003258315.1 | train |
| GCA_002841945.1 | ASM284194v1 | SAMN06767642 | Deltaproteobacteria bacterium HGW-Deltaproteobacteria-14 | GCA_002841945.1 | train |
| GCA_002731275.1 | ASM273127v1 | SAMN07618665 | Deltaproteobacteria bacterium | GCA_002731275.1 | train |
| GCA_002709835.1 | ASM270983v1 | SAMN07618676 | Deltaproteobacteria bacterium | GCA_002709835.1 | train |
| GCA_002296975.1 | ASM229697v1 | SAMN06455911 | Deltaproteobacteria bacterium UBA796 | GCA_002296975.1 | train |
| GCA_002238945.1 | ASM223894v1 | SAMN05959529 | Proteobacteria bacterium bin106 | GCA_002238945.1 | train |
| GCA_002343185.1 | ASM234318v1 | SAMN06453627 | Deltaproteobacteria bacterium UBA2409 | GCA_002343185.1 | train |
| GCA_001799195.1 | ASM179919v1 | SAMN04314405 | Deltaproteobacteria bacterium RIFOXYA12_FULL_61_11 | GCA_001799195.1 | train |
| GCA_001770175.1 | ASM177017v1 | SAMN04314756 | Bdellovibrionales bacterium RIFOXYD1_FULL_55_31 | GCA_001770175.1 | train |
| GCA_002787535.1 | ASM278753v1 | SAMN06659493 | Deltaproteobacteria bacterium CG11_big_fil_rev_8_21_14_0_20_45_16 | GCA_002787535.1 | train |
| GCA_002344565.1 | ASM234456v1 | SAMN06450675 | Bdellovibrionaceae bacterium UBA2351 | GCA_002344565.1 | train |
| GCA_001798105.1 | ASM179810v1 | SAMN04315323 | Deltaproteobacteria bacterium RIFCSPHIGHO2_02_FULL_40_11 | GCA_001798105.1 | train |
| GCA_002342225.1 | ASM234222v1 | SAMN06453347 | Bdellovibrionaceae bacterium UBA2466 | GCA_002342225.1 | train |
| GCA_001798715.1 | ASM179871v1 | SAMN04316063 | Deltaproteobacteria bacterium RIFCSPLOWO2_02_FULL_50_16 | GCA_001798715.1 | train |
| GCA_000307955.1 | ASM30795v1 | SAMN01813472 | Desulfovibrio magneticus str. Maddingley MBC34 | GCA_000307955.1 | train |
| GCA_002387725.1 | ASM238772v1 | SAMN06450526 | Nitrospirae bacterium UBA4572 | GCA_002387725.1 | train |
| GCA_002347965.1 | ASM234796v1 | SAMN06453794 | Nitrospirae bacterium UBA2233 | GCA_002347965.1 | train |
| GCA_000421065.1 | ASM42106v1 | SAMN02440899 | Acidobacteriaceae bacterium TAA166 | GCF_000421065.1 | train |
| GCA_000820845.2 | K22 | SAMEA3158464 | Pyrinomonas methylaliphatogenes | GCF_000820845.2 | train |
| GCA_002451015.1 | ASM245101v1 | SAMN06451230 | Acidobacteria bacterium UBA6911 | GCA_002451015.1 | train |
| GCA_003222295.1 | ASM322229v1 | SAMN08912114 | Acidobacteria bacterium | GCA_003222295.1 | train |
| GCA_003223145.1 | ASM322314v1 | SAMN08912088 | Acidobacteria bacterium | GCA_003223145.1 | train |
| GCA_002731215.1 | ASM273121v1 | SAMN07618220 | Acidobacteria bacterium | GCA_002731215.1 | train |
| GCA_002238705.1 | ASM223870v1 | SAMN05959517 | Acidobacteria bacterium bin61 | GCA_002238705.1 | train |
| GCA_003152095.1 | 20120700_P3D | SAMN08179850 | Acidobacteria bacterium | GCA_003152095.1 | train |
| GCA_002297665.1 | ASM229766v1 | SAMN06452253 | Holophagaceae bacterium UBA692 | GCA_002297665.1 | train |
| GCA_002747255.1 | ASM274725v1 | SAMN07757424 | Acidobacteria bacterium | GCA_002747255.1 | train |
| GCA_002789615.1 | ASM278961v1 | SAMN06660039 | Acidobacteria bacterium CG_4_9_14_3_um_filter_49_7 | GCA_002789615.1 | train |
| GCA_002010725.1 | ASM201072v1 | SAMN06226397 | Candidatus Aminicenantes bacterium JdFR-80 | GCA_002010725.1 | train |
| GCA_002402325.1 | ASM240232v1 | SAMN06453056 | Candidatus Aminicenantes bacterium UBA4820 | GCA_002402325.1 | train |
| GCA_002898535.1 | ASM289853v1 | SAMD00093761 | bacterium HR11 | GCA_002898535.1 | train |
| GCA_001784175.1 | ASM178417v1 | SAMN04315822 | Candidatus Lindowbacteria bacterium RIFCSPLOWO2_12_FULL_62_27 | GCA_001784175.1 | train |
| GCA_002780065.1 | ASM278006v1 | SAMN06659243 | Candidatus Hydrogenedentes bacterium CG07_land_8_20_14_0_80_42_17 | GCA_002780065.1 | train |
| GCA_001804045.1 | ASM180404v1 | SAMN04314093 | Omnitrophica bacterium RBG_13_46_9 | GCA_001804045.1 | train |
| GCA_001804225.1 | ASM180422v1 | SAMN04315226 | Omnitrophica bacterium RIFCSPLOWO2_01_FULL_50_24 | GCA_001804225.1 | train |
| GCA_002085025.1 | ASM208502v1 | SAMN06560474 | Candidatus Omnitrophica bacterium 4484_213 | GCA_002085025.1 | train |
| GCA_001730085.1 | AB1_pvc | SAMN03766366 | PVC group bacterium (ex Bugula neritina AB1) | GCA_001730085.1 | train |
| GCA_003167475.1 | 20110700_P2D | SAMN08178906 | candidate division FCPU426 bacterium | GCA_003167475.1 | train |
| GCA_001778375.1 | ASM177837v1 | SAMN04314657 | Candidatus Firestonebacteria bacterium RIFOXYD2_FULL_39_29 | GCA_001778375.1 | train |
| GCA_003136635.1 | 20110800_S3S | SAMN08179243 | candidate division FCPU426 bacterium | GCA_003136635.1 | train |
| GCA_002428325.1 | ASM242832v1 | SAMN06453266 | bacterium UBP4_UBA6127 | GCA_002428325.1 | train |
| GCA_001873945.1 | ASM187394v1 | SAMN04328251 | Candidatus Desantisbacteria bacterium CG2_30_40_21 | GCA_001873945.1 | train |
| none | none | none | none | GCA_003498085.1 | train |
| none | none | none | none | GCA_003485015.1 | train |
| GCA_002478245.1 | ASM247824v1 | SAMN06455583 | bacterium UBP18_UBA7526 | GCA_002478245.1 | train |
| GCA_001871015.1 | ASM187101v1 | SAMN04328222 | Candidatus Desantisbacteria bacterium CG1_02_38_46 | GCA_001871015.1 | train |
| GCA_002791075.1 | ASM279107v1 | SAMN06660111 | bacterium (Candidatus Ratteibacteria) CG_4_9_14_3_um_filter_41_21 | GCA_002791075.1 | train |
| GCA_000402295.1 | ASM40229v1 | SAMN02441749 | Candidatus Aerophobus profundus SCGC AAA252-L23 | GCA_000402295.1 | train |
| GCA_002311025.1 | ASM231102v1 | SAMN06457421 | Elusimicrobia bacterium UBA1174 | GCA_002311025.1 | train |
| GCA_001871125.1 | ASM187112v1 | SAMN04328241 | Elusimicrobia bacterium CG1_02_37_114 | GCA_001871125.1 | train |
| GCA_002412545.1 | ASM241254v1 | SAMN06455354 | Elusimicrobia bacterium UBA5214 | GCA_002412545.1 | train |
| GCA_002402375.1 | ASM240237v1 | SAMN06454096 | Elusimicrobia bacterium UBA4817 | GCA_002402375.1 | train |
| GCA_003153935.1 | 20120700_E3M | SAMN08179779 | Elusimicrobia bacterium | GCA_003153935.1 | train |
| GCA_900321865.1 | Rumen uncultured genome RUG730 | SAMEA104666744 | uncultured Elusimicrobia bacterium | GCA_900321865.1 | train |
| GCA_002780705.1 | ASM278070v1 | SAMN06659190 | Elusimicrobia bacterium CG03_land_8_20_14_0_80_50_18 | GCA_002780705.1 | train |
| GCA_001563805.1 | ASM156380v1 | SAMN04123323 | Natrialbaceae archaeon B1-Br10_E2g2 | GCA_001563805.1 | train |
| GCA_002506105.1 | ASM250610v1 | SAMN06027537 | Methanolinea sp. UBA275 | GCA_002506105.1 | train |
| GCA_003158275.1 | 20120600_E3D | SAMN08179605 | Euryarchaeota archaeon | GCA_003158275.1 | train |
| GCA_000011005.1 | ASM1100v1 | SAMD00060931 | Methanocella paludicola SANAE | GCA_000011005.1 | train |
| GCA_003194435.1 | ASM319443v1 | SAMN08434988 | ANME-1 cluster archaeon | GCA_003194435.1 | train |
| GCA_000025505.1 | ASM2550v1 | SAMN02598503 | Ferroglobus placidus DSM 10642 | GCF_000025505.1 | train |
| none | none | none | none | UBA8915 | train |
| GCA_002153915.1 | ASM215391v1 | SAMN05770049 | Methanonatronarchaeum thermophilum | GCF_002153915.1 | train |
| GCA_002505695.1 | ASM250569v1 | SAMN06027237 | Euryarchaeota archaeon UBA487 | GCA_002505695.1 | train |
| GCA_001595915.1 | ASM159591v1 | SAMN04495911 | Thermoplasmatales archaeon SG8-52-3 | GCA_001595915.1 | train |
| GCA_003009755.1 | ASM300975v1 | SAMN06020844 | Thermoplasmatales archaeon SW_10_69_26 | GCA_003009755.1 | train |
| GCA_002495235.1 | ASM249523v1 | SAMN06027064 | Euryarchaeota archaeon UBA287 | GCA_002495235.1 | train |
| GCA_001507955.1 | ASM150795v1 | SAMN03445119 | Methanobacteriaceae archaeon 41_258 | GCA_001507955.1 | train |
| GCA_000006175.2 | ASM617v2 | SAMN00000040 | Methanococcus voltae A3 | GCF_000006175.1 | train |
| GCA_002201895.1 | ASM220189v1 | SAMN06246566 | Methanopyrus sp. SNP6 | GCA_002201895.1 | train |
| GCA_001577775.1 | ASM157777v1 | SAMN03323778 | Pyrococcus kukulkanii | GCF_001577775.1 | train |
| GCA_003229935.1 | ASM322993v1 | SAMN09240142 | Euryarchaeota archaeon | GCA_003229935.1 | train |
| GCA_001515205.2 | ASM151520v2 | SAMN04159681 | Hadesarchaea archaeon YNP_45 | GCA_001515205.2 | train |
| GCA_001316245.1 | ASM131624v1 | SAMD00000523 | Vulcanisaeta sp. JCM 14467 | GCA_001316245.1 | train |
| GCA_001717035.1 | ASM171703v1 | SAMN04994459 | Candidatus Methanomethylicus mesodigestum | GCA_001717035.1 | train |
| GCA_000019605.1 | ASM1960v1 | SAMN02598368 | Candidatus Korarchaeum cryptofilum OPF8 | GCF_000019605.1 | train |
| GCA_001563335.1 | ASM156333v1 | SAMN03998759 | Candidatus Thorarchaeota archaeon SMTZ1-45 | GCA_001563335.1 | train |
| GCA_001940725.1 | ASM194072v1 | SAMN04924815 | Candidatus Heimdallarchaeota archaeon LC_2 | GCA_001940725.1 | train |
| GCA_002779595.1 | ASM277959v1 | SAMN06659232 | archaeon CG07_land_8_20_14_0_80_38_8 | GCA_002779595.1 | train |
| GCA_000220375.1 | ASM22037v1 | SAMN01940911 | Candidatus Nanosalina sp. J07AB43 | GCA_000220375.1 | train |
| GCA_002254415.1 | ASM225441v1 | SAMN06647971 | Candidatus Aenigmarchaeota archaeon ex4484_52 | GCA_002254415.1 | train |
| GCA_002789275.1 | ASM278927v1 | SAMN06660022 | archaeon (Candidatus Huberarchaea) CG_4_9_14_0_8_um_filter_31_21 | GCA_002789275.1 | train |
| none | none | none | none | UBA10154 | train |
| GCA_002687935.1 | ASM268793v1 | SAMN07618708 | Euryarchaeota archaeon | GCA_002687935.1 | train |
| GCA_002792915.1 | ASM279291v1 | SAMN06659736 | Candidatus Micrarchaeota archaeon CG_4_10_14_0_2_um_filter_60_11 | GCA_002792915.1 | train |
| none | none | none | none | UBA10219 | train |
| GCA_001723835.1 | ASM172383v1 | SAMN04979201 | Candidatus Altiarchaeales archaeon WOR_SM1_79 | GCA_001723835.1 | train |
| GCA_000328565.1 | ASM32856v1 | SAMN02261384 | Mycobacterium sp. JS623 | GCF_000328565.1 | valid |
| GCA_001613245.1 | ASM161324v1 | SAMD00018778 | Nocardia speluncae NBRC 108251 | GCF_001613245.1 | valid |
| GCA_900101685.1 | IMG-taxon 2651870117 annotated assembly | SAMN05216174 | Alloactinosynnema iranicum | GCF_900101685.1 | valid |
| GCA_000829695.1 | ASM82969v1 | SAMD00020221 | Streptomyces sp. NBRC 110035 | GCF_000829695.1 | valid |
| GCA_900091955.1 | IMG-taxon 2657245721 annotated assembly | SAMN04883155 | Streptomyces sp. Ncost-T10-10d | GCF_900091955.1 | valid |
| GCA_000719865.1 | ASM71986v1 | SAMN02645498 | Streptomyces albus subsp. albus | GCF_000719865.1 | valid |
| GCA_000425985.1 | ASM42598v1 | SAMN02440853 | Microbacterium sp. URHA0036 | GCF_000425985.1 | valid |
| GCA_001685415.1 | ASM168541v1 | SAMN04578256 | Serinicoccus sp. JLT9 | GCF_001685415.1 | valid |
| GCA_900094555.1 | IMG-taxon 2615840609 annotated assembly | SAMN04487811 | Rhizobium hainanense | GCF_900094555.1 | valid |
| GCA_000015165.1 | ASM1516v1 | SAMN02598359 | Bradyrhizobium sp. BTAi1 | GCF_000015165.1 | valid |
| GCA_001043895.1 | ASM104389v1 | SAMN03217758 | Methylobacterium indicum | GCF_001043895.1 | valid |
| GCA_000384335.2 | ASM38433v2 | SAMN02194552 | Hyphomicrobium sp. 99 | GCF_000384335.2 | valid |
| GCA_001464655.1 | Rhizobiales bacterium Ga0077555 | SAMN04254526 | Rhizobiales bacterium Ga0077555 | GCA_001464655.1 | valid |
| GCA_000281715.1 | ASM28171v1 | SAMN00839615 | Sphingobium sp. AP49 | GCF_000281715.1 | valid |
| GCA_002002925.1 | ASM200292v1 | SAMN06263770 | Sphingomonas sp. LM7 | GCF_002002925.1 | valid |
| GCA_000936425.1 | ASM93642v1 | SAMN03382941 | Skermanella aerolata KACC 11604 | GCF_000936425.1 | valid |
| GCA_000300255.2 | ASM30025v2 | SAMN03081445 | Candidatus Methanomethylophilus alvus Mx1201 | GCF_000300255.2 | valid |
| GCA_001481295.1 | ASM148129v1 | SAMN04326629 | Candidatus Methanomethylophilus sp. 1R26 | GCF_001481295.1 | valid |
| GCA_002504495.1 | ASM250449v1 | SAMN06027054 | Methanomassiliicoccaceae archaeon UBA537 | GCA_002504495.1 | valid |
| GCA_002496385.1 | ASM249638v1 | SAMN06027036 | Methanomassiliicoccales archaeon UBA147 | GCA_002496385.1 | valid |
| GCA_002505185.1 | ASM250518v1 | SAMN06027152 | Ferroplasmaceae archaeon UBA567 | GCA_002505185.1 | valid |
| GCA_002498845.1 | ASM249884v1 | SAMN06027607 | Thermoplasmatales archaeon UBA509 | GCA_002498845.1 | valid |
| GCA_002254385.1 | ASM225438v1 | SAMN06647974 | Thermoplasmatales archaeon ex4484_6 | GCA_002254385.1 | valid |
| GCA_002254765.1 | ASM225476v1 | SAMN06647967 | Thermoplasmatales archaeon ex4484_36 | GCA_002254765.1 | valid |
| GCA_003151735.1 | 20120700_P3D | SAMN08179876 | Candidatus Bathyarchaeota archaeon | GCA_003151735.1 | valid |
| GCA_001273335.1 | ASM127333v1 | SAMN03793026 | miscellaneous Crenarchaeota group-1 archaeon SG8-32-3 | GCA_001273335.1 | valid |
| GCA_002010975.1 | ASM201097v1 | SAMN06226324 | Candidatus Bathyarchaeota archaeon JdFR-07 | GCA_002010975.1 | valid |
| GCA_001399795.1 | ASM139979v1 | SAMN03983875 | Candidatus Bathyarchaeota archaeon BA2 | GCA_001399795.1 | valid |
| GCA_002490245.1 | ASM249024v1 | SAMN07701645 | Candidatus Bathyarchaeota archaeon B24-2 | GCA_002490245.1 | valid |
| GCA_001776015.1 | ASM177601v1 | SAMN04316258 | Candidatus Bathyarchaeota archaeon RBG_16_57_9 | GCA_001776015.1 | valid |
| GCA_001915065.1 | ASM191506v1 | SAMN04327914 | archaeon 13_2_20CM_2_53_6 | GCA_001915065.1 | valid |
| GCA_003152955.1 | 20120700_E3M | SAMN08179789 | Candidatus Bathyarchaeota archaeon | GCA_003152955.1 | valid |
| GCA_002479715.1 | ASM247971v1 | SAMN06454495 | Ruminococcus sp. UBA7477 | GCA_002479715.1 | test |
| GCA_003312465.1 | ASM331246v1 | SAMN08494074 | Faecalibacterium prausnitzii | GCA_003312465.1 | test |
| GCA_002315275.1 | ASM231527v1 | SAMN06456561 | Clostridiales bacterium UBA1777 | GCA_002315275.1 | test |
| none | none | none | none | GCA_003522145.1 | test |
| GCA_002308775.1 | ASM230877v1 | SAMN06456139 | Mageeibacillus sp. UBA1257 | GCA_002308775.1 | test |
| GCA_001916035.1 | ASM191603v1 | SAMN05717248 | Clostridium sp. 27_14 | GCA_001916035.1 | test |
| GCA_900066385.1 | 13414_6#69 | SAMEA3545293 | uncultured Clostridium sp. | GCA_900066385.1 | test |
| GCA_002841495.1 | ASM284149v1 | SAMN06767700 | Firmicutes bacterium HGW-Firmicutes-2 | GCA_002841495.1 | test |
| GCA_000436595.1 | MGS1124 | SAMEA3138714 | Prevotella sp. CAG:1124 | GCA_000436595.1 | test |
| GCA_900316005.1 | Rumen uncultured genome RUG152 | SAMEA104666332 | uncultured Bacteroidales bacterium | GCA_900316005.1 | test |
| GCA_002429385.1 | ASM242938v1 | SAMN06451136 | Prolixibacteraceae bacterium UBA6029 | GCA_002429385.1 | test |
| GCA_003259835.1 | ASM325983v1 | SAMN06297150 | Flavobacterium aquaticum | GCF_003259835.1 | test |
| GCA_001425355.1 | Leaf405 | SAMN04151757 | Chryseobacterium sp. Leaf405 | GCF_001425355.1 | test |
| GCA_002690915.1 | ASM269091v1 | SAMN07618573 | Crocinitomicaceae bacterium | GCA_002690915.1 | test |
| GCA_003054115.1 | ASM305411v1 | SAMN08777206 | Pedobacter sp. YR016 | GCF_003054115.1 | test |
| GCA_900102545.1 | IMG-taxon 2634166296 annotated assembly | SAMN04488121 | Chitinophaga filiformis | GCF_900102545.1 | test |
| GCA_001714685.2 | ASM171468v2 | SAMN05438611 | Methanosarcina sp. Ant1 | GCA_001714685.2 | test |
| GCA_000970085.1 | ASM97008v1 | SAMN03074589 | Methanosarcina siciliae T4/M | GCF_000970085.1 | test |
| GCA_900114835.1 | IMG-taxon 2642422540 annotated assembly | SAMN04488696 | Methanolobus profundi | GCF_900114835.1 | test |
| GCA_000970325.1 | ASM97032v1 | SAMN03074571 | Methanococcoides methylutens MM1 | GCF_000970325.1 | test |
| GCA_002926195.1 | ASM292619v1 | SAMN06562579 | ANME-2 cluster archaeon HR1 | GCA_002926195.1 | test |
| none | none | none | none | UBA10536 | test |
| GCA_002501765.1 | ASM250176v1 | SAMN06027569 | Methanosaeta sp. UBA204 | GCA_002501765.1 | test |
| GCA_003162615.1 | 20100900_E3D | SAMN08178753 | Methanothrix sp. | GCA_003162615.1 | test |
| GCA_000484975.1 | ASM48497v1 | SAMN02440765 | Thaumarchaeota archaeon SCGC AAA282-K18 | GCA_000484975.1 | test |
| GCA_003175215.1 | ASM317521v1 | SAMD00113975 | Candidatus Nitrosopumilus sp. NM25 | GCF_003175215.1 | test |
| GCA_000484935.1 | ASM48493v1 | SAMN02441105 | Euryarchaeota archaeum SCGC AAA287-E17 | GCA_000484935.1 | test |
| GCA_001510295.1 | ASM151029v1 | SAMN03733546 | Thaumarchaeota archaeon casp-thauma4 | GCA_001510295.1 | test |
| GCA_900143675.1 | Candidatus Nitrosotalea sinensis Nd2 whole genome | SAMEA20449918 | Nitrosotalea sp. Nd2 | GCF_900143675.1 | test |
| GCA_002713205.1 | ASM271320v1 | SAMN07620155 | Thaumarchaeota archaeon | GCA_002713205.1 | test |
| GCA_002011085.1 | ASM201108v1 | SAMN06226331 | Candidatus Geothermarchaeota archaeon JdFR-14 | GCA_002011085.1 | test |
| GCA_002898355.1 | ASM289835v1 | SAMD00093751 | archaeon HR01 | GCA_002898355.1 | test |

**SI Table 1** – NCBI accessions for the genomes included in the *GTDB class set* (Methods).

| Model | bs | bptt | wd | moms | Dropout rates | | | | | optim | loss fn |
|---|---|---|---|---|---|---|---|---|---|---|---|
| | | | | | input | embed | hidden | weight | output | | |
| LookingGlass | 512 | 100 | 1e-2 | (0.98,0.9) | 0.04 | 0.005 | 0.03 | 0.05 | 0.04 | Adam ß=(0.9,0.99) | Cross Entropy |
| Functional annotation classifier | 512 | - | 1e-2 | (0.8,0.7) | 0.12 | 0.015 | 0.09 | 0.15 | 0.12 | Adam ß=(0.9,0.99) | Cross Entropy |
| Oxidoreductase classifier | 256 | - | 1e-2 | (0.8,.07) | 0.12 | 0.015 | 0.09 | 0.15 | 0.12 | Adam ß=(0.9,0.99) | Cross Entropy |
| Translation frame classifier | 256 | - | 1e-2 | (0.8,0.7) | 0.12 | 0.015 | 0.09 | 0.15 | 0.12 | Adam ß=(0.9,0.99) | Cross Entropy |
| Optimal temperature classifier | 256 | - | 1e-2 | (0.8,0.7) | 0.12 | 0.015 | 0.09 | 0.15 | 0.12 | Adam ß=(0.9,0.99) | Cross Entropy |

**SI Table 2** – Hyperparameter settings for the LookingGlass language model and the fine tuned transfer learning classifiers. Column name abbreviations stand for: bs=batch size; bptt=back propagation through time; wd=weight decay; moms=momentums; the variable dropout rates determine the frequency of dropout applied to the dropout mask in each section of the model: input=frequency of dropout in the dropout mask for the data input matrix; embed=dropout frequency for the embedding matrix; hidden=dropout frequency for the hidden layers; weight=dropout frequency for the hidden layer weights; output=dropout frequency for the encoder output layer.

| Environmental package | SRA Run Accession |
|---|---|
| built environment | ERR1332600 |
| built environment | SRR1577782 |
| built environment | ERR1332623 |
| built environment | ERR1332614 |
| built environment | ERR1332606 |
| built environment | ERR1332586 |
| built environment | SRR1577774 |
| built environment | ERR2699805 |
| built environment | ERR2699809 |
| host-associated | SRR3184383 |
| host-associated | SRR925829 |
| host-associated | ERR1366724 |
| host-associated | ERR1135232 |
| host-associated | ERR2241851 |
| host-associated | ERR2200504 |
| host-associated | ERR2200674 |
| host-associated | ERR2765138 |
| human-gut | ERR209626 |
| human-gut | SRR5091465 |
| human-gut | ERR1600429 |
| human-gut | ERR209703 |
| human-gut | SRR5164026 |
| human-gut | ERR1190830 |
| human-gut | ERR3053395 |
| human-gut | ERR2816294 |
| human-gut | ERR3521978 |
| microbial mat/biofilm | SRR1171647 |
| microbial mat/biofilm | SRR1707409 |
| microbial mat/biofilm | ERR1855538 |
| microbial mat/biofilm | ERR1855552 |
| microbial mat/biofilm | SRR2556884 |
| microbial mat/biofilm | ERR1855546 |
| microbial mat/biofilm | ERR1739691 |
| microbial mat/biofilm | SRR830624 |
| microbial mat/biofilm | ERR2020026 |

| | |
|---|---|
| microbial mat/biofilm | ERR2020015 |
| miscellaneous | SRR1298754 |
| miscellaneous | DRR046818 |
| miscellaneous | DRR027592 |
| miscellaneous | ERR1698988 |
| miscellaneous | ERR1353140 |
| miscellaneous | ERR2239851 |
| miscellaneous | ERR2298558 |
| miscellaneous | ERR1358726 |
| miscellaneous | ERR2239841 |
| miscellaneous | ERR2239840 |
| plant-associated | SRR1511001 |
| plant-associated | SRR1754159 |
| plant-associated | ERR2145397 |
| plant-associated | ERR2145413 |
| plant-associated | ERR2709726 |
| plant-associated | ERR2969987 |
| plant-associated | ERR2819892 |
| plant-associated | ERR2709750 |
| plant-associated | ERR2144884 |
| plant-associated | ERR2022395 |
| sediment | SRR4069407 |
| sediment | ERR1474613 |
| sediment | ERR1560098 |
| sediment | ERR970605 |
| sediment | ERR1201181 |
| sediment | ERR1743388 |
| sediment | ERR1743304 |
| sediment | ERR1743339 |
| sediment | ERR2215874 |
| sediment | ERR1743283 |
| soil | SRR1574704 |
| soil | ERR1017187 |
| soil | SRR1238204 |
| soil | ERR1700691 |
| soil | SRR1238205 |
| soil | SRR5234512 |
| soil | ERR2603191 |
| soil | ERR1877921 |
| soil | ERR1960504 |
| soil | ERR2767288 |
| wastewater/sludge | ERR712383 |
| wastewater/sludge | SRR2938315 |
| wastewater/sludge | ERR977414 |
| wastewater/sludge | ERR977422 |
| wastewater/sludge | ERR1076075 |
| wastewater/sludge | SRR1616983 |
| wastewater/sludge | ERR1076080 |
| wastewater/sludge | ERR1960627 |
| wastewater/sludge | ERR1746303 |
| wastewater/sludge | ERR3173383 |
| water | SRR4343439 |
| water | ERR598955 |
| water | ERR1726775 |
| water | ERR1726943 |
| water | ERR1726572 |
| water | ERR694158 |
| water | ERR599252 |
| water | SRR1185414 |
| water | ERR1987930 |
| water | ERR2017141 |

**SI Table 3** – SRA IDs and their associated environmental packages for environmental metagenomes used for the creation of the *mi-faser functional set* (Methods).

| SRA read accession | Lat | Lon | Ocean region | Depth category | Depth (m) | Temp (°C) | Oxygen (μmol/kg) | OMZ? | TARA Station |
|---|---|---|---|---|---|---|---|---|---|
| ERR598981 | -12.93 | -96.12 | Pacific | MES | 175.3 | 13.0 | 0.7 | Yes | 100 |
| ERR599063 | -12.99 | -95.99 | Pacific | SRF | 5.5 | 25.3 | 200.2 | - | 100 |
| ERR599115 | 35.31 | -127.74 | Pacific | MES | 644.6 | 4.9 | 8.6 | Yes | 133 |
| ERR599052 | 35.41 | -127.74 | Pacific | SRF | 5.5 | 19.2 | 224.4 | - | 133 |
| ERR599020 | -1.87 | -84.62 | Pacific | MES | 376.9 | 10.3 | 1.3 | Yes | 110 |
| ERR599039 | -2.01 | -84.59 | Pacific | SRF | 5.5 | 23.9 | 190.8 | - | 110 |
| ERR599076 | 14.17 | -116.66 | Pacific | MES | 371.0 | 8.9 | 0.6 | Yes | 137 |
| ERR598989 | 14.20 | -116.63 | Pacific | SRF | 5.4 | 26.4 | 195.1 | - | 137 |
| ERR599048 | -8.80 | -17.91 | Atlantic | MES | 792.7 | 4.7 | 143.2 | No | 072 |
| ERR599105 | -8.78 | -17.91 | Atlantic | SRF | 5.8 | 25.0 | 199.1 | - | 072 |
| ERR598964 | 34.10 | -49.78 | Atlantic | MES | 734.3 | 10.6 | 155.2 | No | 149 |
| ERR598963 | 34.10 | -49.89 | Atlantic | SRF | 5.5 | 18.7 | 220.2 | - | 149 |
| ERR599125 | -61.98 | -49.45 | Polar | MES | 783.8 | 0.5 | 203.8 | No | 085 |
| ERR599176 | -62.03 | -49.54 | Polar | SRF | 5.9 | 0.7 | 343.4 | - | 085 |
| ERR3589593 | 76.12 | 1.36 | Polar | MES | 490.5 | 0.1 | 315.4 | No | 163 |
| ERR3589586 | 76.18 | 1.39 | Polar | SRF | 5.0 | 1.9 | 363.0 | - | 163 |

**SI Table 4** – SRA read accessions and associated metadata for metagenomes in the *oxidoreductase metagenome set* (Methods).

| Taxlevel | Max accuracy | Max accuracy cutoff | >90% precision cutoff | >95% precision cutoff | >98% precision cutoff | $R^2$ emb vs. seq similarity | % pairs <50 seq similarity |
|---|---|---|---|---|---|---|---|
| **Genus** | 0.789 | 0.7 | 0.82 | 0.88 | 0.91 | 0.44 | 10.7% |
| **Family** | 0.766 | 0.67 | 0.84 | 0.89 | 0.92 | 0.41 | 17.3% |
| **Order** | 0.732 | 0.63 | 0.84 | 0.89 | 0.92 | 0.37 | 23.2% |
| **Class** | 0.683 | 0.61 | 0.88 | 0.91 | 0.93 | 0.30 | 45.0% |
| **Phylum** | 0.664 | 0.62 | 0.89 | 0.92 | 0.94 | 0.28 | 44.4% |

**SI Table 5** – Metrics for differentiation between homologous and nonhomologous sequence pairs for the five levels of taxonomic specificity tested. Columns: 'Max accuracy'=maximum accuracy across all cutoffs; 'Max accuracy cutoff'=cutoff at which maximum accuracy is achieved; '>90%, 95%, 98% precision cutoff'=cutoff at which at least 90%, 95%, or 98% precision is achieved; '$R^2$ emb vs. seq similarity'=correlation coefficient between embedding cosine similarity and sequence similarity bit scores for each comparison; '% pairs <50 seq similarity'=percent of homologous sequence comparisons for which the sequence similarity bit score is less than 50.

| SRA read accession | Oxidoreductase classifier | MG-RAST | | mi-faser | |
|---|---|---|---|---|---|
| | % oxidoreductases | % annotated | % oxido | % annotated | % oxido |
| ERR599105 | 16.40 | 27.90 | 2.32 | 1.61 | 0.38 |
| ERR599063 | 18.18 | 40.31 | 3.69 | 0.17 | 0.04 |
| ERR598989 | 18.21 | 37.61 | 3.63 | 2.41 | 0.48 |
| ERR598963 | 18.30 | 28.96 | 2.94 | 1.83 | 0.34 |
| ERR599039 | 18.46 | - | - | 2.34 | 0.46 |
| ERR3589593 | 18.47 | 29.04 | 2.69 | 1.82 | 0.29 |
| ERR599048 | 18.53 | 30.67 | 2.75 | 1.88 | 0.39 |
| ERR599052 | 18.59 | 36.86 | 3.58 | 1.27 | 0.25 |
| ERR599176 | 18.62 | 50.26 | 4.03 | 2.72 | 0.47 |
| ERR599020 | 18.74 | 30.13 | 3.07 | 1.94 | 0.33 |
| ERR599125 | 19.23 | 26.70 | 2.22 | 1.51 | 0.25 |
| ERR3589586 | 19.64 | 47.19 | 0.02 | 2.86 | 0.59 |
| ERR598964 | 19.75 | 30.33 | 2.92 | 1.73 | 0.28 |
| ERR599115 | 20.01 | 31.75 | 2.79 | 1.01 | 0.16 |
| ERR598981 | 20.23 | 33.16 | 3.20 | 2.17 | 0.34 |
| ERR599076 | 20.61 | 32.06 | 0.01 | 1.88 | 0.29 |

**SI Table 6** – Comparison of the % oxidoreductases predicted by the oxidoreductase classifier relative to the MG-RAST and mi-faser functional annotation tools for the metagenomes in the *oxidoreductase metagenome set,* as well as the % reads annotated overall for MG-RAST and mi-faser.

#### References for Supporting Online Material


1. Parks, D. H. *et al.* A standardized bacterial taxonomy based on genome phylogeny substantially revises the tree of life. *Nat. Biotechnol.* **36**, 996 (2018).
